# Lineage-specific X chromosome inactivation escape and skew underlie sex-biased immune gene dosage and deleterious variant exposure

**DOI:** 10.64898/2026.08.26.739472

**Authors:** Daisy Kavanagh, Alice Steel, Helen E. King, Helaine Graziele Santos Vieira, Kishore R. Kumar, Etienne Masle-Farquhar, Cecile King, Robert Weatheritt, Ksenia Skvortsova

**Author notes:** These authors contributed equally (Robert Weatheritt, Ksenia Skvortsova).

## Abstract

The X chromosome carries an unusually high density of immune genes and is a major contributor to sex differences in immune function and autoimmune diseases. In females, X-chromosome inactivation (XCI) has two major functional consequences: it shapes X-linked gene dosage through XCI escape and determines the cellular exposure of heterozygous X-linked variants through XCI skew. Yet because XCI creates a mosaic of cells expressing different parental X chromosomes, these properties have remained largely inaccessible in individual women, becoming measurable only where XCI is non-random or after aggregation across large cohorts. Consequently, how X-linked variation contributes to sex-biased immunity and differs between individual women has remained unresolved. Here we present scDaisyChain, a graph-based framework that reconstructs chromosome-scale X haplotypes directly from heterozygous SNPs and single-cell long-read transcriptomes. scDaisyChain achieves near-ground-truth accuracy in highly polymorphic mouse hybrids and shows strong concordance with orthogonal long-read whole-genome phasing in human samples. Applied to peripheral blood immune cells from healthy women, it reveals a lineage-specific escape program in which lymphoid cells escape XCI more broadly than monocytes, with corresponding gains in the inactive X chromatin accessibility and female-biased expression. Lineage-specific skew further alters the proportion of cells expressing each heterozygous X-linked variant, a property we term variant exposure. Predicted deleterious variants are preferentially found in low-exposure states, exemplified by a splice-altering *TLR8* variant expressed in few cytotoxic T cells. In rheumatoid arthritis (RA), the monocyte compartment - which has the lowest escape in health - shows reproducible inactive X dysregulation converging on a trained-immunity programme linked to disease flare and synovial macrophage activation, with elevated escape of *IL13RA1* and *HDAC8*. These findings establish lineage-specific escape, skew and variant exposure as quantifiable, patient-resolved determinants of sex-biased immune gene dosage and X-linked variant penetrance in health and autoimmune disease, resolving a dimension of female biology that has been previously inaccessible in individual donors.

**One Sentence Summary:** scDaisyChain resolves X-inactivation escape, skew and variant exposure in individual women, shaping immune dosage and autoimmune risk.

## INTRODUCTION

The X chromosome harbours genes essential for immune function and neurodevelopment and is increasingly recognised as a major contributor to sex-biased tissue function and disease susceptibility^1–3^. Despite this importance, the X-chromosome has historically been underrepresented in large-scale genomic studies^4–5^, obscuring our understanding of its contribution to phenotypic variation between sexes and among females. A major source of this variation arises from X chromosome inactivation (XCI), an epigenetic process that stochastically silences one of the two X chromosomes in XX female cells to achieve dosage compensation with XY males^6^. Foundational work has established long non-coding RNA (lncRNA) *XIST* as the master regulator of XCI, silencing the second X chromosome through the formation of supramolecular condensates, recruiting RNA binding proteins, histone deacetylases and Polycomb to drive repressive chromatin remodelling, compaction and stable gene silencing^7–12^ - consistent with the broader capacity of lncRNAs to scaffold chromatin-modifying complexes at specific genomic loci^13^.

Despite the breadth of *XIST*-mediated silencing, approximately 15-30% of X-linked genes in humans escape silencing and are expressed from both the active (Xa) and inactive (Xi) X chromosomes^14–16^. Escape from XCI is highly heterogeneous: some genes consistently escape inactivation, while others show partial or context-dependent escape varying across tissues, cell types, and developmental stages^16^. This variability creates substantial differences in X-linked gene dosage between individuals and cellular populations. Accumulating evidence indicates that allele-specific X-linked gene expression, and particularly escape from XCI, contributes to sex-biased differences in immune function^1–2,17–20^, neurological diseases^3,21–22^, autoimmune disorders^3,19,23–26^ and responses to infection^27^. This growing recognition that X chromosome activity influences clinically relevant phenotypes, has renewed interest in comprehensive approaches for profiling XCI across tissues, cell types, and disease states^16,28–30^.

Despite this growing recognition of the biological importance of XCI escape, its systematic characterisation in human cells remains challenging. A key obstacle is the stochastic nature of XCI, as most female tissues contain a heterogeneous mix of cells differing in which parental allele is inactivated, making bulk samples poorly suited to quantify XCI escape. This limitation has been circumvented by using model systems: human cell lines or somatic-cell hybrid systems with clonal or highly skewed XCI^15^, rare primary human samples with naturally skewed XCI^16,31^ and mouse models in which XCI can be genetically controlled^32–34^. However, each approach has limitations. Immortalised human cell lines frequently harbour chromosomal abnormalities and may not accurately reflect normal physiological contexts, while bulk RNA-seq in mouse models cannot readily resolve cell type variability of XCI, and the extreme levels of XCI skew required to perform this analysis with bulk RNA sequencing is rare in humans. As a result, the extent and consequences of XCI escape across human cell types and disease states remains incompletely understood.

Single-cell long-read sequencing substantially alleviates the limitations of short-read approaches by sequencing transcript-spanning reads, substantially increasing the probability that a read will overlap an informative heterozygous SNP. However, exploiting this additional allelic information requires chromosome-scale phasing of X linked variants. Existing phasing approaches remain poorly suited to this task: (i) read-based phasing is fragmented by sparse variant density; (ii) trio-based phasing requires parental genotypes that are often unavailable; (iii) population-based approaches are restricted to common variants and remain susceptible to switch errors; (iv) and methylation-based approaches cannot readily be applied to the X chromosome, because methylation predominantly reflects XCI status rather than parental haplotype.

To overcome these limitations, we developed scDaisyChain, a graph-based framework for chromosome-scale reconstruction of X haplotypes from single-cell long-read transcriptomes. Without requiring parental genotypes, highly skewed XCI or cohort-level aggregation, scDaisyChain enables direct quantification of XCI escape, XCI skew and variant exposure within individual donors. We show that lineage-specific variation in X chromosome inactivation shapes immune gene dosage, redistributes the cellular exposure of heterozygous X-linked variants and is perturbed in female-biased autoimmune disease.

## RESULTS

### Chromosome-scale reconstruction of X haplotypes enables patient-resolved analysis of X chromosome inactivation

Single-cell analyses of X chromosome inactivation (XCI) require assignment of transcripts to the active and inactive X chromosomes (Xa and Xi). In human samples, this remains challenging because heterozygous SNPs are sparse and conventional short-read sequencing captures only a small fraction of each transcript sequence, leaving many X-linked genes without an informative SNP overlap. Although long-read single-cell sequencing substantially increases the probability of detecting heterozygous variants, no framework currently reconstructs chromosome-scale X haplotypes or enables direct quantification of XCI escape, XCI skew and variant exposure within individual donors.

To address this challenge, we developed scDaisyChain, a graph-based framework that reconstructs chromosome-scale X haplotypes from expressed heterozygous variants and quantifies XCI activity directly from single-cell long-read transcriptomes (Fig. 1). scDaisyChain exploits the principle that alleles residing on the same parental chromosome are repeatedly co-expressed within individual cells. A community detection algorithm first identifies a high-confidence scaffold of phased heterozygous variants from allelic co-expression patterns, which is iteratively refined before chromosome-scale phase is extended to the remaining variants by comparing their cell-specific allelic expression with haplotype-specific expression profiles (Fig 1a, b). This reconstructs chromosome-scale haplotypes directly from single-cell transcriptomic data without requiring parental genotypes or cohort-level aggregation.

**Figure 1.**
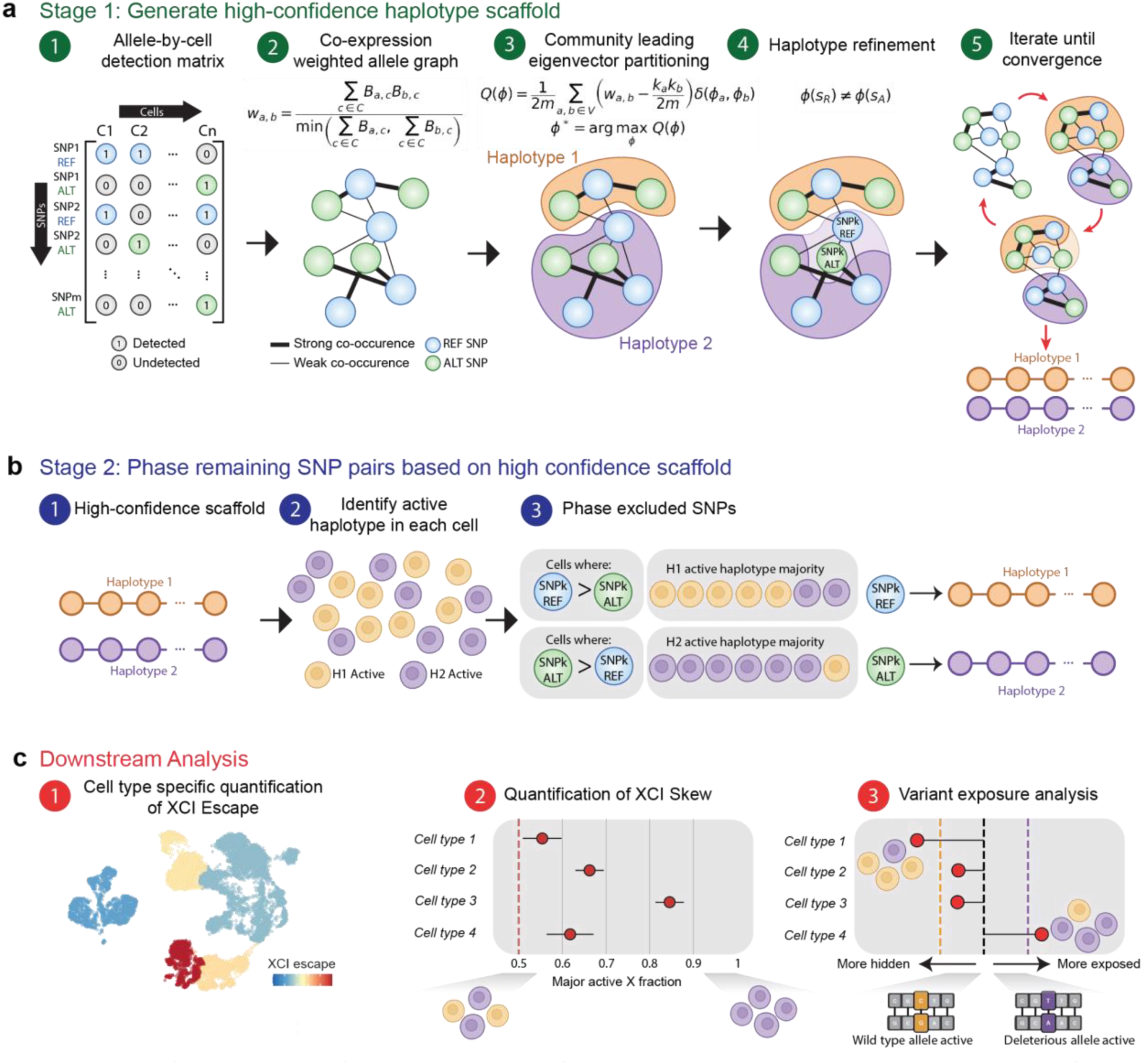
Overview of the scDaisyChain phasing algorithm. scDaisyChain reconstructs the parental X chromosome haplotypes and quantifies expression from the active and inactive X. **(a)** In Stage 1, heterozygous SNPs are encoded in a binary detection matrix, which is then used to construct a weighted graph where the edges represent co-expression frequency of a pair of alleles. Community leading eigenvector partitioning is used to separate the graph into two broad haplotypes, which are then further refined by removal of pairs of alleles where the REF and ALT segregate into the same haplotype. The graphing, partitioning and refinement are repeated iteratively until there is convergence to two haplotypes with no further refinement possible. **(b)** In stage 2, pairs of SNPs excluded from stage 1 are phased based on the concordance of their allelic expression with the active X identity of each cell. **(c)** The final haplotypes enable downstream analyses including cell type specific quantification of XCI escape, estimation of XCI skew and variant exposure analysis.

The reconstructed haplotypes are subsequently used to assign single-cell reads to the active and inactive X chromosomes, generating allele-specific expression matrices that enable direct quantification of XCI escape, cell-type-specific XCI skew and heterozygous variant exposure, providing a unified framework for patient-resolved X chromosome biology analysis (Fig. 1c).

### Accurate chromosome-scale X phasing in mouse and human single-cell data

We next benchmarked scDaisyChain using datasets with known haplotypes and orthogonal measures of X-chromosome activity to assess the accuracy, robustness and biological validity of the framework.

We first used an established XCI model, female mouse F1 hybrids from a C57BL/6 x CAST/EiJ cross, which is a widely used allele-specific model due to the high number of polymorphisms between strains, and provides chromosome-scale ground-truth haplotypes^35–37^. Applying scDaisyChain to this dataset, the initial graph-based phasing step assigned 3,982 of 4,769 expressed X-linked SNPs with 99.97% accuracy, while the remaining variants were then assigned in a second stage using haplotype-informed cell-level expression patterns, resulting in an overall phasing accuracy of 98.09% across all expressed SNPs (Fig. 2a). Residual SNP-level errors had minimal impact on downstream biological inference. Aggregation of phased allelic counts across genes and cells generated expression profiles that showed extremely strong agreement with the ground-truth strain assignment at both gene and cell levels (Pearson’s r = 0.998–1.000, Supplementary Materials Fig. S1a). Importantly, scDaisyChain produces ratios of Xi expression per cell and per gene that are very strongly correlated with the ground truth (Pearson’s r = 0.97–0.986, Fig. 2b,d). Even genes that contain one or more misclassified SNPs produce Xi ratios that are comparable to the ground truth, highlighting the robustness of our model (Supplementary Materials Fig. S1b). Overall, most genes show no deviation in their Xi ratio from the ground truth, with only 4 genes showing a deviation of 0.05 or greater (Supplementary Materials Fig. S1c,d). These results demonstrate that scDaisyChain accurately reconstructs biologically meaningful chromosome-scale haplotypes despite sparse single-cell allelic data.

**Figure 2.**
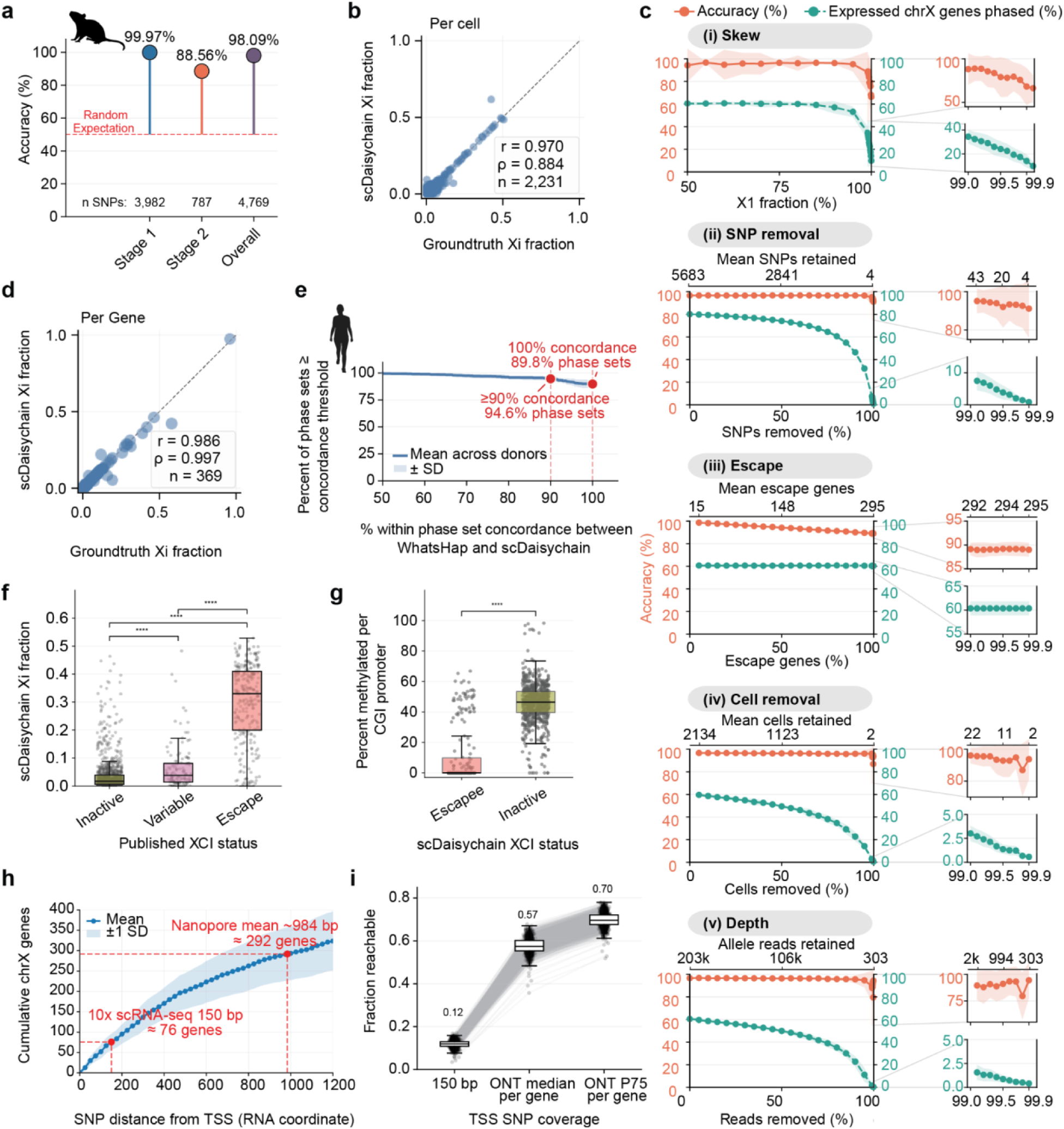
Benchmarking scDaisyChain for chromosome-scale haplotype reconstruction and XCI quantification. **(a)** Accuracy per stage of the scDaisyChain algorithm applied to the C57BL/6 X CAST/EiJ F1 mouse data with known ground truth. **(b)** Fraction of X chromosome expression from the inactive X per cell aggregated across all genes in the ground truth mouse data vs results obtained from scDaisyChain. **(c)** Simulated single-cell allelic expression data using the C57/Bl6 X CAST/EiJ mouse data as the starting point and varying (i) XCI skew, (ii) Number of heterozygous SNPs, (iii) Percentage of genes that escape XCI, (iv) Number of cells in the dataset, (v) Allelic read depth. For all simulations, XCI skew was set to 50-50, 80% SNP removal, 25% escape, all cells, unless it was the parameter being tested. **(d)** Fraction of X chromosome expression from the inactive X per gene aggregated across all cells. **(e)** Concordance of within phase set phasing from WhatsHap of Nanopore WGS of human samples and scDaisyChain results. For each phase set identified by WhatsHap, the phasing of each SNP was compared to the scDaisyChain phasing result. SNPs were considered concordant if the phasing result matched between the two methods after the best phase set orientation. **(f)** scDaisyChain Xi fraction of genes from human data, and their previous classification as inactive, variable or escape from Tukiainen et al., 2017^16^. **(g)** CpG island methylation at promoters of genes by scDaisyChain classified XCI status. **(h)** Average number of genes with a heterozygous X chromosome SNP in female individuals in the 1kGenomes project at a range of distances from the transcriptional start site (TSS), with typical 10x Illumina read length and mean read length in our 10x Nanopore human samples. **(i)** Average proportion of PBMC expressed X chromosome genes per female individual in the 1K genomes project with a SNP reachable in the first 150 exonic bp, using the median or 75th percentile read length of the gene from our 10x Nanopore human samples.

Although the C57BL/6 x CAST/EiJ mouse F1 benchmark provides chromosome-scale ground truth, human samples typically exhibit lower heterozygosity, more balanced XCI and a greater proportion of escaping genes. To assess performance under these conditions, we generated simulated datasets by systematically varying XCI skew, escape frequency, and SNP density (Fig. 2c). Across a broad range of parameter settings, scDaisyChain maintained high phasing accuracy, with appreciable declines observed only under extreme scenarios such as >99% XCI skew or near-complete escape across the X chromosome. In the case of XCI skew, increased skew results in an increased proportion of SNPs phased in stage 2 of the model compared to stage 1, however with this comes an increase in stage 2 phasing accuracy leading to the overall phasing accuracy remaining stable (Supplementary Materials Fig. S1f). Reductions in SNP density primarily reduced the number of classifiable genes rather than phasing accuracy itself, indicating that the framework is robust to the sparse allelic information characteristic of human single-cell datasets. We further simulated technical variation including the number of cells and read depth, and observed a similar pattern to SNP removal: primarily a reduction in the number of classifiable genes while accuracy was maintained until extreme levels of removal (>99% removal; Fig. 2c). Collectively, these simulations indicate that the principal consequence of reduced information content is loss of classifiable genes rather than loss of phasing accuracy.

We next sought to directly validate scDaisyChain phasing in human samples using an orthogonal long-read DNA-based approach. To obtain a reference set of heterozygous variants and their phasing independent of RNA expression, we performed whole-genome nanopore sequencing and phased X-chromosome variants using WhatsHap^38^. Because read-based DNA phasing is constrained by variant density and read connectivity, it typically yields multiple phase sets rather than a single chromosome-scale haplotype. We therefore compared scDaisyChain and WhatsHap within individual phase blocks, where relative haplotype assignment is known. Within these phase sets, scDaisyChain showed strong concordance with WhatsHap-derived phasing results with 94.6% of phase sets showing ≥ 90% concordance between the two methods (Fig. 2e), with an average concordance per phase set of 98.2% (Supplementary Materials Fig. S2a). Discordance was concentrated within the pseudoautosomal region (PAR1) and regions proximal to XIST, where constitutive escape results in near-balanced allelic expression, providing limited information for haplotype assignment (Supplementary Materials Fig. S2b). Importantly, correcting the within phase set haplotype assignment to match the WhatsHap derived results had minimal impact on the Xi ratio per gene for the vast majority of genes, with strong correlations between the estimates from both methods (Supplementary Materials Fig. S2c).

We next assessed whether scDaisyChain-derived Xi estimates in human data recapitulated established features of X-chromosome inactivation biology. Consistent with known XCI status, genes previously classified as escapees exhibited substantially higher Xi fractions than variably escaping or fully inactivated genes (Fig. 2f). Using a threshold of 10% Xi-derived expression to define escape, scDaisyChain classifications showed strong agreement with published XCI annotation, with only 18/328 genes (∼5.49%) showing discordant escape / inactive classification (Supplementary Materials Fig. S2h). These discrepancies may reflect biological differences between tissues, cell types or donor cohorts. In addition, scDaisyChain assigned XCI status to 27 genes that had not been previously classified. Consistent with these observations, using nanopore whole-genome sequencing data, we observed that genes classified as escapees showed significantly reduced promoter CpG island and CpG island shore methylation at promoter regions compared with inactivated genes (Fig. 2g, Supplementary Materials Fig. S2d-g), providing an orthogonal epigenetic validation of scDaisyChain-derived escape estimates. These benchmarks demonstrate that scDaisyChain reconstructs chromosome-scale X haplotypes with high accuracy across species and under human-like conditions, providing a validated foundation for allele-specific analyses in individual donors.

### Long-read transcript coverage enables per-donor XCI phasing

Patient-resolved XCI analysis requires sufficient allelic information per donor to phase X-linked genes without pooling across individuals. We therefore asked how single-cell long-read sequencing changes the fraction of the X chromosome that can be interrogated within an individual donor. Using female genomes from the 1000 Genomes Project, we quantified the distribution of heterozygous exonic SNP positions across X-linked transcripts and estimated the number of potentially classifiable genes as a function of accessible transcript length (Fig. 2h)^39–40^. Restricting analysis to a 150bp window downstream of the TSS, approximating conventional 10x short-read 5’ capture, yielded a median of 76 informative genes per individual. In contrast, extending the accessible transcript window to the mean nanopore read length in our dataset (984 nucleotides) increased this to 292 genes, representing a 3.84-fold increase in potential XCI-resolved gene recovery. However, this single global read-length threshold provides a conservative summary of the gain from long-read sequencing, because observed read span varies substantially between genes, and is constrained by the transcript architecture, because genes with genuinely short transcript lengths will pull down the global mean. To better capture this variability, we calculated the median read length for each expressed gene in our human PBMC dataset. The average female individual in the 1000 Genomes project had a reachable heterozygous SNP in only 12% of these genes under 10x-equivalent 150 bp coverage, compared to 57% using gene-specific median nanopore read spans, further increasing to 70% using the 75th percentile read span. (Fig. 2i). Similar increases in classifiable genes with longer nanopore read lengths were observed when counting from the transcriptional end site, mimicking 10x 3’ capture (Supplementary Materials Fig. S3a,b). To confirm that these theoretical gains translated to allelic information in practice, we truncated our nanopore reads to 150 bp on average, which reduced the number of SNP-overlapping reads per cell to ∼14.7% of that seen with full-length reads (a 6.8-fold gain from long reads; Supplementary Materials Fig. S3c). These results suggest that transcript-spanning reads substantially expand the fraction of the X-chromosome that can be interrogated within a single individual.

Together, long-read transcript coverage substantially increases the number of X-linked genes amenable to allele-specific analysis, enabling patient-resolved XCI interrogation that would not be feasible with short-read data.

### scDaisyChain detects cell type-specific XCI escape in healthy immune cells

To demonstrate scDaisyChain’s utility across biological contexts, we first examined XCI escape in healthy human immune cells. We isolated ∼2,000-4,000 peripheral blood mononuclear cells (PBMCs) from four healthy women and performed single-cell capture followed by ONT long-read RNA-seq (Supplementary Table 1). Parallel ONT whole-genome sequencing identified germline heterozygous SNPs distinguishing the two X chromosomes. To account for the sparsity inherent in allele-resolved single-cell measurements, Xi fractions were regularised using empirical beta-prior shrinkage. These regularised Xi fractions were used for all downstream analyses, providing a quantitative, donor-aware framework for evaluating Xi fraction across immune cell types. Using scDaisyChain, we quantified the fraction of expression from the inactive X chromosome (Xi fraction) for informative SNP-containing X-linked genes in single-cells across major immune cell types, revealing notable variation in escape across cell types (Fig. 3a, b, Supplementary Materials Fig. S3a,b). To identify genes with cell type-variable Xi escape, we classified X-linked genes into PAR, constitutive escape, variable escape, and inactive/unknown categories based on the Tukiainen *et al.* classification^16^ and calculated the mean and standard deviation of Xi fraction across cell types for each gene. As expected, most PAR genes and several known escapees showed constitutive biallelic expression across all PBMC cell types (Fig. 3c, bottom right). Consistent with these classifications, mean Xi fractions showed strong agreement with independent escape estimates derived from scLinaX, particularly for constitutive and variable escape genes (Supplementary Materials Fig. S3c)^30^. However, a substantial subset of genes displayed cell type-variable Xi escape (Fig. 3c, top left), including both known escape genes and X-linked genes not previously annotated as escapees (Supplementary Materials Fig. S3d).

**Figure 3.**
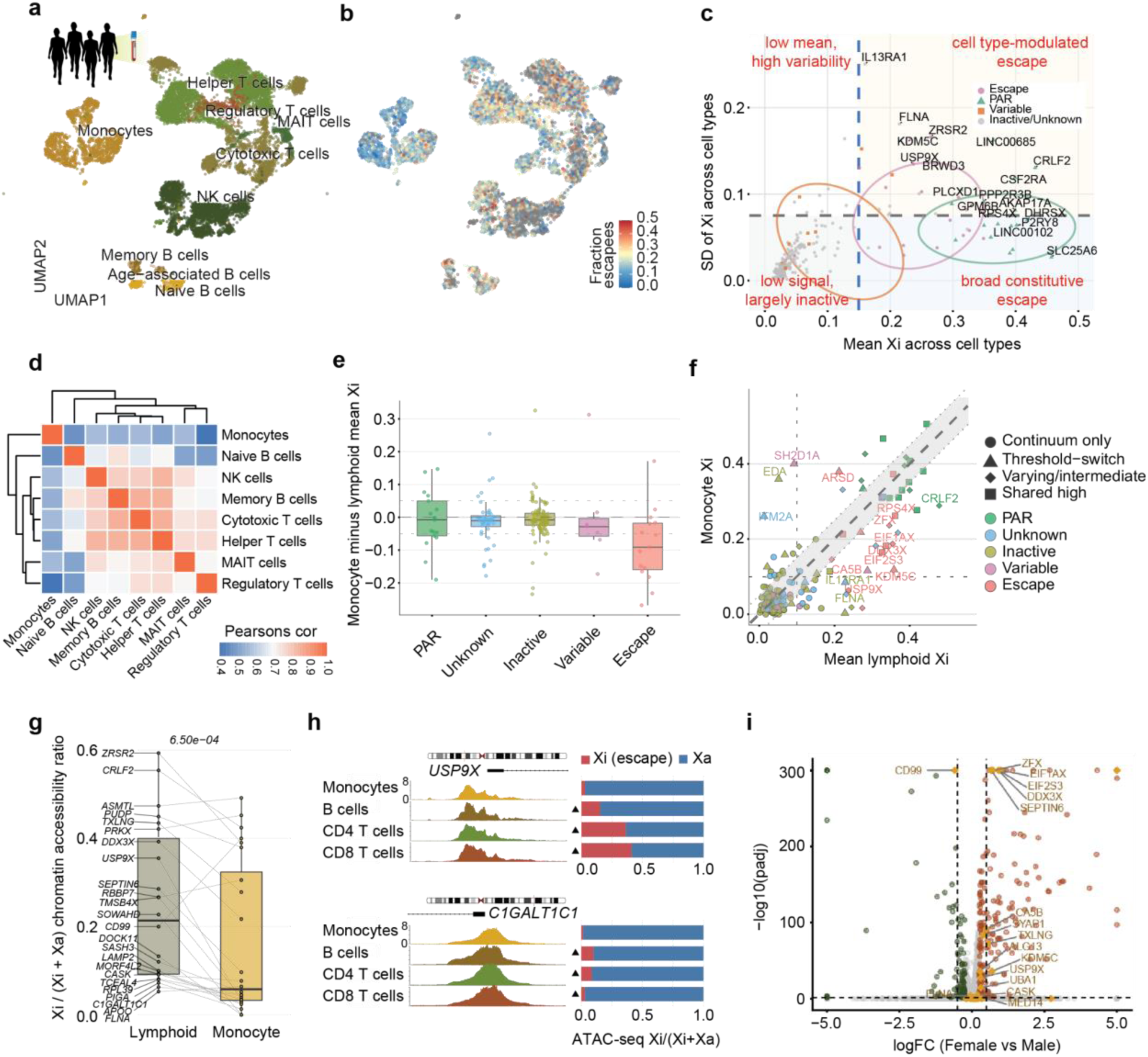
Cell type-specific escape from X chromosome inactivation in female immune cells. **(a)** UMAP projection of healthy female PBMCs coloured by cell type annotation. **(b)** UMAP projection coloured by fraction of genes escaping XCI per cell, showing heterogeneous escape burden across cell types. **(c)** Mean Xi fraction versus standard deviation across cell types for each X-linked gene. Genes in the upper right quadrant exhibit cell type-modulated escape, while genes in the lower right show broad constitutive escape. Genes with low mean and high variability (upper left) represent loci exhibiting cell type-restricted escape. Point shape denotes XCI annotation class. **(d)** Pairwise Pearson correlation of Xi escape profiles across cell types. Monocytes cluster separately from the lymphoid compartment, indicating distinct escape programmes. **(e)** Distribution of monocyte minus lymphoid mean Xi fraction per gene, stratified by XCI annotation class. Known escape genes show the greatest monocyte–lymphoid divergence. **(f)** Monocyte versus lymphoid mean Xi fraction for individual genes. Points above the diagonal indicate higher Xi fraction in monocytes; points below indicate higher Xi in lymphoid cells. Point colour denotes XCI annotation class and point shape indicates the escape mode. **(g)** Xi-derived chromatin accessibility ratio (scLinaX-multi, Tomofuji et al. 2024) for lymphoid escapee genes in lymphoid versus monocyte compartments, with paired lines connecting the same gene. Lymphoid escapees show higher Xi accessibility in lymphoid cells (p = 8.5 × 10⁻⁴). **(h)** IGV browser tracks and stacked barplots showing ATAC-seq profiles (left) and Xi/(Xi+Xa) chromatin accessibility ratio across cell types for two representative genes: *USP9X* and *C1GALT1C1*, illustrating cell type-specific chromatin opening on the inactive X. **(i)** Volcano plot of sex-biased gene expression (female versus male) across lymphoid cells from the Allen Immune Health Atlas. Lymphoid escape genes identified by scDaisyChain (yellow diamonds) are enriched among female-upregulated X-linked genes, confirming that Xi escape translates into sex-biased dosage.

We next examined the overlap of escaping genes across cell types (Supplementary Materials Fig. S3e). The largest group (I1, n = 17) consisted of genes escaping in all cell types, comprising predominantly known constitutive escapees and PAR genes, confirming robust detection of allele-specific escape at single-cell resolution across donors. By contrast, genes with escape restricted to specific cell types (I2–I12) were predominantly classified as variable escapees, indicating that cell-type-specific escape is shaped by context-dependent rather than constitutive regulatory mechanisms.

To further explore cell type-specific Xi escape profiles, we calculated pairwise Pearson correlations using donor-aggregated mean Xi fractions for non-PAR X-linked genes (Fig. 3d). Most cell-type pairs showed high correlation (r > 0.7), indicating a shared core Xi escape programme. The notable exception was monocytes, whose Xi escape profiles were consistently less similar to those of lymphoid populations (r ∼ 0.5) (Fig. 3d). Monocytes also exhibited the lowest mean Xi fraction (14.5%) of any immune cell type (Supplementary Materials Fig. S3f), suggesting that monocytes differ from lymphoid cells in both the pattern and overall level of Xi escape. To investigate the basis for this divergence between myeloid and lymphoid Xi escape profiles, we grouped cell types by developmental lineage and calculated gene-level differences in Xi fraction (Fig. 3e). PAR genes showed no significant difference between lineages (p = 0.294), consistent with their biallelic expression being independent of XCI. In contrast, known constitutive escapees displayed significantly lower Xi fraction in monocytes compared to lymphoid cells (p < 0.001), indicating that the myeloid–lymphoid divergence in Xi escape is primarily driven by reduced escape of constitutive escapee genes in the myeloid lineage. Genes with lymphoid-specific escape were enriched for immune signalling and pathogen sensing functions (Fig. 3f), including immune signal transduction (*DDX3X*, *USP9X*), and complement regulation (*CFP*), as well as chromatin regulators with known roles in immune cell differentiation (*KDM5C*) and were enriched for cytokine signalling, receptor activity and immune-associated pathways (Supplementary Materials Fig. S3g). Using publicly available allele-resolved single-cell ATAC-seq^30^, we confirmed that promoters of lymphoid-specific escapees had higher chromatin accessibility on the Xi in lymphocytes but not in monocytes, consistent with their cell type-specific Xi escape status (Fig. 3g, h). More broadly, constitutive escapees had the highest chromatin accessibility on the Xi, followed by cell type-specific escapees, while promoters of inactive genes had closed chromatin on the Xi (Supplementary Materials Fig. S3h,i), highlighting the robustness of escape identification by scDaisyChain.

Finally, we asked whether Xi escape results in elevated expression of X-linked genes in female compared to male immune cells. Differential gene expression analysis of the Allen Human Immune Cell Atlas^41^ (dataset DOI: 10.57785/e9e1-wh09) revealed that genes escaping XCI in the lymphoid lineage had significantly higher expression in female B cells compared to males (Fig. 3i and Supplementary Materials Fig. S3j), including genes involved in innate immune signalling and interferon responses (*DDX3X*, *USP9X*)^42–43^, immune cell homeostasis (*ZFX*)^44^, and chromatin regulation (*KDM5C*)^45^. These results suggest that lymphoid-specific Xi escape underlies sex-biased gene dosage of key immune regulators.

Taken together, these results show that scDaisyChain enables robust identification of cell type-specific escape from XCI at single-donor resolution. The observed patterns suggest that Xi escape is shaped in part by developmental lineage, with a major divergence between myeloid and lymphoid programmes, and that this escape translates into sex-biased dosage of immune genes, providing a potential molecular basis for sex disparities in immunity and autoimmune disease susceptibility.

### Rheumatoid arthritis is associated with monocyte-specific Xi dysregulation

Having established a healthy reference dataset of cell-type-specific XCI escape in healthy female PBMCs, we next investigated whether Xi fraction is altered in autoimmune disease (AID). AID display a pronounced female bias, yet the contribution of cell-type-specific XCI dysregulation to disease susceptibility remains poorly understood. To address this, we applied long-read single-cell RNA sequencing and scDaisyChain to PBMCs from female patients with rheumatoid arthritis (RA; n = 3) - an AID with a 3:1 female-to-male ratio^46^ - and compared Xi-fraction profiles to those of healthy female controls.

We first quantified disease-associated changes in Xi fraction across immune cell populations. Projection of the average RA-healthy difference in escape fraction onto the PBMC atlas revealed marked heterogeneity across cell types (Fig. 4a-b, Supplementary Materials Fig. S4a). Rather than a uniform increase in escape across immune cell types, alterations were concentrated within specific compartments: monocytes exhibited the largest increase in Xi fraction relative to healthy controls, while memory B cells and helper T cells showed more modest increases, and most other cell types displayed little evidence of systematic change (Fig. 4b and Supplementary Materials Fig. S4b-c). Permutation testing confirmed that monocytes exhibited the strongest evidence for increased escape fraction among all cell populations examined (Fig. 4c; p < 0.015).

**Figure 4.**
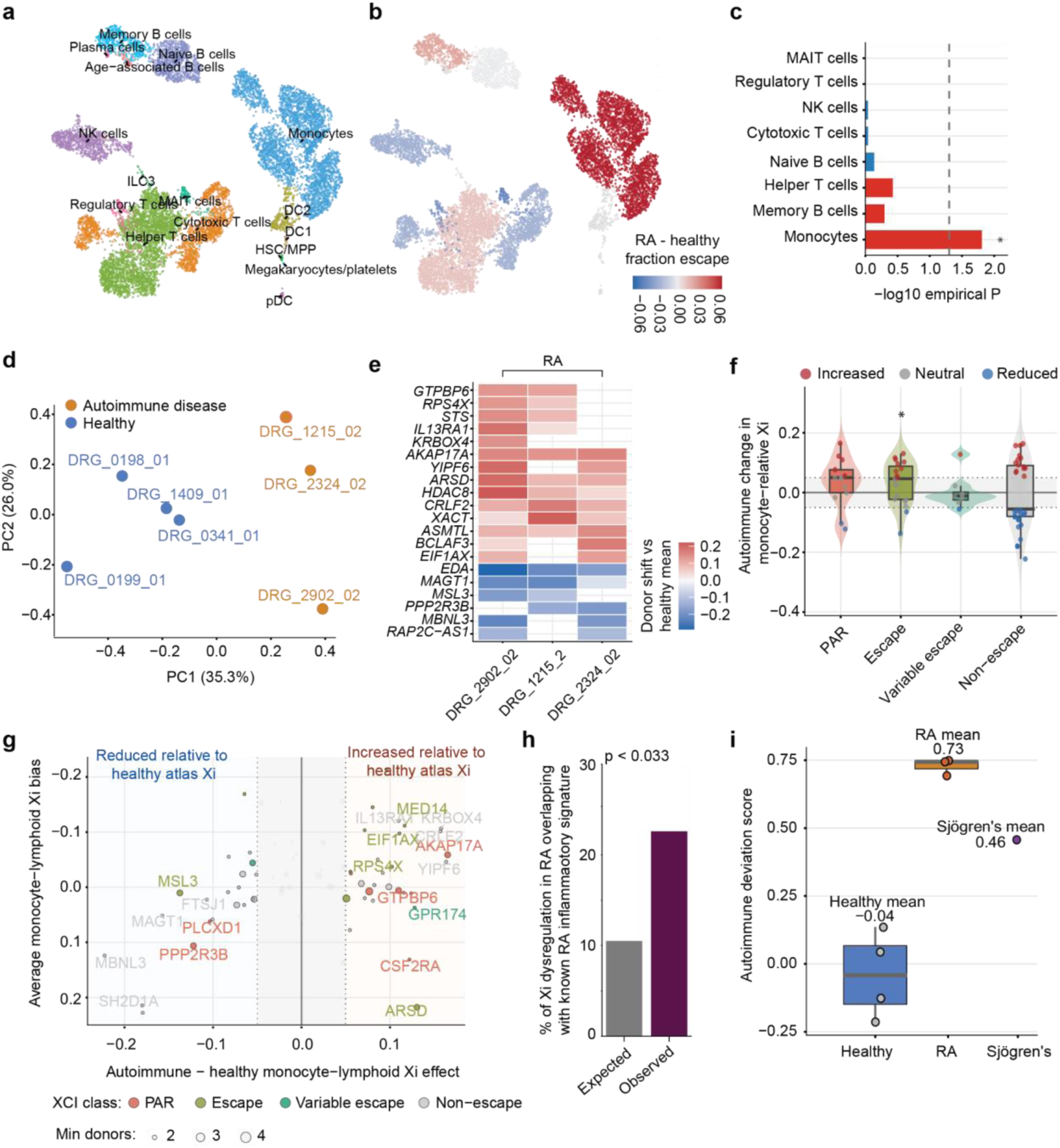
Cell type-specific escape from X chromosome inactivation in autoimmune disease. **(a)** UMAP projection of PBMCs from female rheumatoid arthritis patients coloured by cell type annotation. **(b)** UMAP projection coloured by differences in fraction of genes escaping XCI per cell between healthy and rheumatoid arthritis patients. **(c)** Cell-type-specific permutation analysis assessing evidence for net escape-associated deviation relative to the healthy reference atlas. Monocytes exhibited the strongest enrichment for coordinated autoimmune-associated Xi escape alterations. * = p < 0.05. **(d)** Principal component analysis (PCA) plot of monocyte-specific Xi fraction with each dot representing a separate donor. Fraction of variance for each axis is displayed in brackets. PC = principal component **(e)** Heatmap showing donor-level Xi shifts relative to the healthy mean for representative X-linked genes across RA donors, demonstrating reproducible directional changes across individuals. **(f)** Boxplot and violin plots showing autoimmune-associated changes in monocytes relative to the healthy controls. Escape associated genes showed preferentially positive shifts in Xi escape in RA donors, whereas non-escape genes showed minimal directional bias. **(g)** Gene-level autoimmune-associated Xi effects plotted against baseline monocyte–lymphoid Xi bias. Genes positioned in the right quadrant exhibited increased relative Xi within monocyte-associated contexts compared with the healthy atlas, whereas genes in the left quadrant showed reduced monocyte-relative Xi. Point colour denotes XCI/PAR annotation class and point size indicates the number of contributing donors. **(h)** Barplot showing overlap between genes exhibiting autoimmune-associated Xi dysregulation and a published monocyte trained-immunity signature associated with RA disease flare and synovial macrophage activation. **(i)** A boxplot showing autoimmune deviation score derived from the RA cohort and applied to an independent female patient with primary Sjögren’s syndrome.

Given the prominence of the monocyte signal, we next performed principal component analysis using monocyte-specific Xi values. Although analysis of all cell types together did not clearly separate healthy and RA donors (Supplementary Materials Fig. S4d), monocyte Xi profiles alone distinguished RA patients from controls along the principal axis of variation (PC1 = 35.3%, PC2 = 26.0%; Fig. 4d and Supplementary Materials Fig. S4e), indicating that disease-associated Xi dysregulation is concentrated within this lineage rather than reflecting a global PBMC-wide effect. To assess the reproducibility of gene-level Xi dysregulation, we examined monocyte Xi shifts in each RA donor relative to the healthy reference mean. Although the magnitude of effect varied across patients, the direction of change was highly concordant across donors with multiple genes exhibiting consistent increases or decreases in Xi fraction in all three individuals (Fig. 4e and Supplementary Materials Fig. S4f). This concordance indicates that the observed Xi signature is not driven by a single patient but reflects a shared pattern of X-linked dosage perturbation across the RA cohort.

We next asked whether disease-associated Xi dysregulation preferentially affected particular classes of X-linked genes. Genes classified as constitutive escapees showed significantly larger autoimmune disease-associated changes than variably escaping or non-escape genes (Fig. 4f; p < 0.0031, Wilcoxon rank-sum test), suggesting that loci already predisposed to incomplete silencing are especially susceptible to further dosage perturbation in disease. Consistent with this, genes exhibiting increased Xi fraction in RA were enriched among established escape genes, whereas genes with reduced Xi fraction were more frequently observed among non-escape loci (Fig. 4g). Furthermore, leave-one-donor-out analyses showed that the monocyte Xi signature remained highly stable when any individual RA donor was excluded, indicating that the observed pattern was not driven by a single patient (Supplementary Materials Fig. S4g). To determine whether genes showing consistent Xi fraction changes across RA patients were linked to established RA biology, we compared them against a published monocyte trained-immunity signature associated with disease flare and synovial macrophage activation^47^. Among the 31 expressed X-linked genes in this signature, 7 exhibited a reproducible RA-associated Xi dysregulation, representing a 2.16-fold observed over expected enrichment (Fig. 4h; p < 0.033, Fisher’s exact test). This overlap provides independent support that reproducible Xi alterations in RA monocytes occur within gene networks previously implicated in inflammatory disease activity.

A key promise of patient-resolved XCI profiling is detection of XCI abnormalities in individuals for whom cohort-scale analysis is not feasible. To test this in an independent disease context, we asked whether an RA-derived deviation score could flag abnormal Xi activity in a single patient with primary Sjögren’s syndrome. The Sjögren’s sample showed a substantially elevated deviation score relative to healthy controls and clustered more closely with RA samples than with healthy controls (Fig. 4i). This observation exceeded all randomised RA-axis projections tested (Supplementary Materials Fig. S4h) and remained robust across alternative gene sets, including exclusion of PAR genes and restriction to escape-relevant non-PAR genes (Supplementary Materials Fig. S4i-j).

Although this single case is insufficient to infer shared disease mechanisms between rheumatoid arthritis and Sjögren’s syndrome, it demonstrates that scDaisyChain-derived Xi profiles can be interpreted at the level of individual patients, without requiring extreme chromosome-wide skewing or large disease cohorts, opening a practical route for studying XCI in rare diseases and precision medicine settings.

### Cell-type-specific XCI skew reshapes the cellular exposure of X-linked variants across immune lineages

A key advantage of scDaisyChain is that chromosome-scale phasing within individual donors enables direct measurement of XCI skew and the resulting cellular exposure of heterozygous X-linked variants. While XCI escape influences X-linked gene dosage, XCI skew determines which haplotype remains active within a given cell population. By shifting the proportion of cells expressing each parental X chromosome, skew directly modulates the cellular exposure of heterozygous X-linked variants across cell types. We therefore quantified XCI skew across donors and immune cell populations to determine how variation in XCI skew reshaped the cellular exposure of heterozygous X-linked variants.

XCI skew varied substantially between individuals, with the major X chromosome active in approximately 50% to over 70% of cells depending on the donor (Fig. 5a, Supplementary Materials Fig. S5a). Within individuals, skew magnitude was further modulated by lineage identity: cytotoxic and innate-like lymphoid populations consistently exhibited greater skew than the donor average, whereas B cell and myeloid populations exhibited reduced skew (Fig. 5b, *P* < 0.039, BH-corrected Wilcoxon test). Together, these findings show that XCI skew is shaped by both donor-specific factors and immune lineage identity, generating systematic variation in the fraction of cells expressing each X haplotype - even within the same individual.

**Figure 5.**
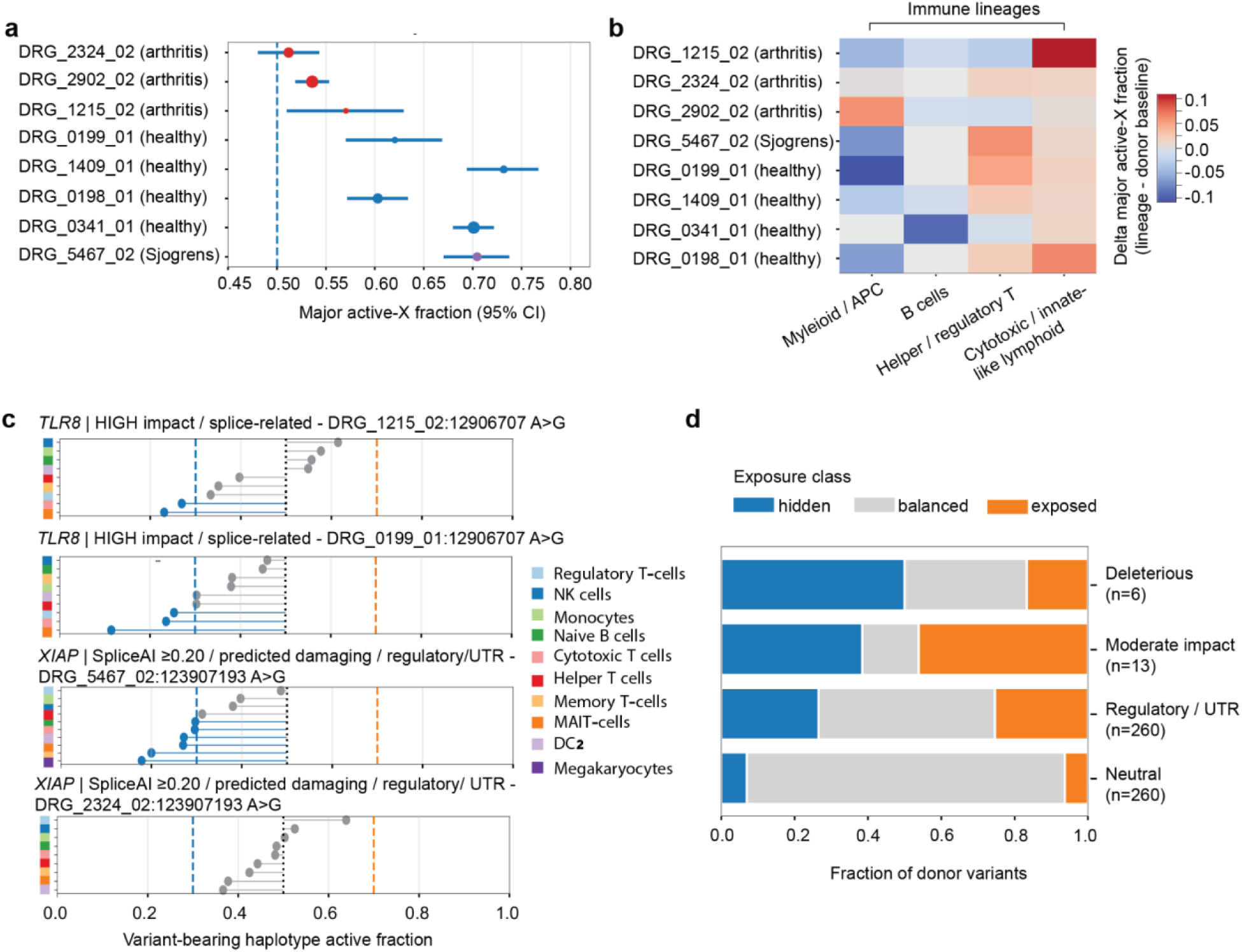
Cell type-specific skew reshapes exposure of heterozygous X-linked variants. **(a)** Global donor-level XCI skew estimated from phased single-cell allelic expression. Points indicate the major-active-X fraction for each donor and error bars represent 95% confidence intervals. The dashed line indicates balanced XCI (0.5). Donors are coloured by cohort. **(b)** Immune lineage-specific deviations in major-active-X fraction relative to each donor’s global baseline. Positive values indicate increased skew within a lineage, whereas negative values indicate reduced skew relative to the donor-wide average. **(c)** Representative examples illustrating lineage-specific variation in variant exposure. Each point indicates the fraction of cells within a given immune population in which the variant-bearing haplotype was active. Blue and orange dashed lines denote hidden (0.3) and exposed (0.7) thresholds, respectively, and the black dotted line indicates balanced exposure (0.5). **(d)** Distribution of donor-variants classified as hidden (variant-bearing haplotype active in ≤30% of cells), balanced, or exposed (variant-bearing haplotype active in ≥70% of cells) across functional annotation categories. Significant enrichment relative to neutral low-priority variants following Benjamini–Hochberg correction of Fisher’s exact tests.

We reasoned that lineage-specific differences in XCI skew should directly alter the cellular exposure of heterozygous X-linked variants by shifting which X chromosome haplotype remains active across cells. To test this, we assigned each phased X-linked variant to its parental haplotype and quantified, for each donor and immune cell type, the proportion of cells in which the variant-bearing haplotype was on the active X. Variants were then classified as hidden (active in fewer than 30% of cells), exposed (active in more than 70% of cells), or balanced (intermediate frequencies).

This analysis revealed that the same heterozygous variant frequently occupied distinct exposure states in immune lineages within the same individual. Consequently, genotype alone was insufficient to predict variant exposure, as lineage-specific XCI skew substantially reshaped the cellular exposure of the same heterozygous X-linked allele (Fig. 5c and Supplementary Materials Fig. S5b). For example, a splice-related *TLR8* variant exhibited markedly reduced cellular exposure in cytotoxic T cells relative to other immune populations from the same donor (Fig. 5c). Together, these findings show that chromosome-scale phasing enables direct quantification of variant exposure as a cell-type-specific consequence of lineage-specific XCI skew.

Next we asked whether exposure patterns differed according to predicted functional consequences. Deleterious variants (VEP high-impact and splice-related variants) were significantly enriched in hidden states relative to neutral variants (odds ratio = 13.3, BH-adjusted q-value = 0.012) with no corresponding enrichment observed among exposed states (Fig. 5d). Although based on a limited number of variants, these findings suggest that variants predicted to have severe functional consequences are more frequently associated with reduced cellular exposure. In contrast, moderate-impact variants and regulatory/UTR variants were observed in both hidden and exposed states, indicating that XCI skew broadly redistributes variant exposure across immune lineages without a consistent directional bias (Fig. 5d and Supplementary Materials Fig. S5c, d).

Together, these analyses show that cell-type-specific XCI skew alters the cellular exposure of heterozygous X-linked variation across immune lineages. Whereas moderate-impact and regulatory variants were redistributed across both hidden and exposed states, deleterious variants were preferentially represented among hidden states, reducing the fraction of cells in which they were expressed. More broadly, these analyses introduce variant exposure as a quantifiable cell-type specific phenotype of X-linked variation, demonstrating that the cellular exposure of a heterozygous X-linked variant depends not only on genotype, but also on the immune lineage in which it is measured.

## DISCUSSION

Systematic characterisation of allele-specific X chromosome activity in females has depended on systems that circumvent cellular heterogeneity rather than resolve it: clonal cell lines, rare donors with naturally skewed inactivation, and mouse crosses with known, highly polymorphic haplotypes or genetically controlled XCI skewing. scDaisyChain removes that dependency by reconstructing chromosome-scale X haplotypes directly from single-cell long-read transcriptomes, enabling XCI escape, XCI skew, and the resulting exposure of heterozygous variants to be quantified within individual donors.

Applied to peripheral blood from healthy donors and women with autoimmune disease, scDaisyChain generated allele-resolved maps of X chromosome activity across immune cell types. Previous studies have shown that escape varies across tissues and individuals^15,33^ and, more recently, across cell types using single-cell RNA sequencing^30^. Our data resolve Xi escape to the level of the individual donor. In healthy individuals, the most striking divergence separated monocytes from lymphoid populations, with the majority of established escape genes exhibiting lower Xi fraction in the myeloid compartment. These findings suggest that XCI escape is not solely an intrinsic property of individual genes but is also shaped by cellular context. Lymphoid-specific escapees showed corresponding increases in Xi chromatin accessibility, implicating lineage-specific chromatin environments in regulating escape^30^. This is consistent with smFISH and immunofluorescence evidence that lncRNA *XIST* is unstable at certain stages of B and T lymphocyte development, accompanied by the depletion of heterochromatin mark H2AK119Ub^23,51–52^, suggesting that the lymphoid-specific escape patterns we observe are modulated by dynamic *XIST* occupancy between cell lineages^53^.

Patient-resolved quantification of X chromosome activity opens a route to investigating X chromosome dysregulation in disease, a long-standing hypothesis for the female bias of autoimmune conditions involving altered escape, skewing and *XIST*-dependent XCI maintenance^18,26,54–57^. Two lines of experimental evidence directly support this model. *XIST* deletion in patient B cells *ex vivo* drove the expansion of atypical B cells, a subset known to accumulate during chronic antigen stimulation in viral infections and autoimmune disease^58^. Independently, *TLR7*, an X-linked innate immune receptor, illustrates how even modest escape from XCI can contribute to pathology: a mild increase in *TLR7* dosage was sufficient to trigger SLE-like systemic autoimmune disease in mice, characterised by anti-RNA autoantibody production and a pro-inflammatory interferon signature^59^. Together, these findings suggest that Xi dysregulation - whether through loss of *XIST*-dependent silencing or escape of specific immunoregulatory loci - may converge to amplify female-biased autoimmune pathology.

In rheumatoid arthritis, the same monocyte compartment that shows the most restrained escape in health is the site of Xi-derived alterations that are reproducible across donors and largely absent from other immune populations. This is notable given the central role of monocytes as precursors of the synovial macrophages and osteoclasts that drive joint inflammation and bone erosion in autoimmune disease^60–61^. Among genes showing increased Xi-derived expression in female RA monocytes, *IL13RA1* exerts pleiotropic immunological functions including suppression of pro-inflammatory cytokine production and regulation of autoreactive B cell expansion^62–64^. Another example is *HDAC8*, a histone deacetylase whose inhibition suppresses osteoclastogenesis in preclinical models^65^, raising the possibility that biallelic expression lowers the epigenetic barrier to osteoclast differentiation and contributes to the female bias in RA-associated bone destruction. Notably, an autoimmune deviation score derived from RA samples identified abnormal Xi activity in an independent patient with Sjögren’s syndrome. These findings demonstrate that disease-associated X-chromosome phenotypes can therefore be quantified in individual patients - a capability of particular value in rare diseases, small cohorts, and historically understudied populations where sufficient sample sizes for population-level XCI analysis are difficult to achieve. Although larger cohorts will be required to determine the prevalence and significance of these alterations, our findings establish a framework for patient-resolved analysis of X chromosome activity across human disease.

XCI skew determines which parental X remains active in each cell and therefore which heterozygous X-linked variants are expressed. XCI skew has long been recognised as a modifier of X-linked disease penetrance^66–67^, but variant exposure has not been quantifiable directly across cell types within individual donors. The consequences of altered variant exposure are perhaps most clearly illustrated in X-linked primary immunodeficiencies including chronic granulomatous disease, agammaglobulinemia and common variable immunodeficiency, where heterozygous carriers manifest disease because skewing preferentially silences the wild-type allele, rendering affected cells functionally hemizygous for the pathogenic variant^66,68–72^. Competition between clones expressing different parental X chromosomes can further shape this exposure: in mice carrying a loss-of-function *STAG2* variant, cells expressing the mutant haplotype were selectively depleted from the lymphoid compartment through continuous competition with wild-type clones^73–74^. Our findings extend this to naturally occurring XCI skew, showing that variant exposure is redistributed across immune populations in healthy donors. Among the donor variants observed in this cohort, predicted-deleterious variants were preferentially classified as hidden relative to neutral variants, though exposure was dependent on the immune lineage. The mechanism underlying this enrichment remains unclear, but these observations suggest that XCI skew acts as a modifier of the cellular exposure of pathogenic alleles in healthy individuals.

Several technical considerations and opportunities for future extension are worth noting. As with any phasing approach applied to human samples, reliable inference of allele-specific expression remains contingent on sufficient heterozygosity, transcriptional activity of genes bearing heterozygous germline variants, and sequencing depth. Although long-read sequencing substantially increases the number of informative genes, not all X-linked loci were classifiable in every donor or cell type. The autoimmune analyses focused on small, deeply sequenced cohorts by design: patient-resolved chromosome-scale phasing at cell-type resolution requires sequencing depth per donor that constrains cohort size in current long-read single-cell workflows. The datasets generated here, to our knowledge, are the deepest single-cell nanopore transcriptomes in human peripheral blood to date, establish that clinically relevant XCI phenotypes are detectable in individual patients and provide a resource for follow-up cohort-scale studies as long-read single-cell sequencing scales. Additionally, the framework was developed for nanopore long-read sequencing, but the underlying graph-based phasing strategy is platform-agnostic and could in principle be applied to other single-cell transcriptomic datasets that provide sufficient transcript coverage and allelic information. Full-length short-read approaches such as Smart-seq2 and Smart-seq3^75–76^ have enabled per-donor XCI studies at high transcript coverage though at significantly lower cell numbers than droplet-based methods. Finally, current XCI annotations are almost exclusively gene-centric. By combining chromosome-scale phasing with full-length transcript sequencing, scDaisyChain provides a potential framework for resolving X chromosome activity at the isoform level, including whether individual splice isoforms differ in their escape status or exhibit haplotype-biased expression. Such analyses may reveal additional layers of dosage regulation that remain obscured by gene-level measurements.

As long-read single-cell sequencing scales, joint measurement of haplotypes, isoforms and cell states will progressively expand the range of allele-specific regulation accessible to direct measurement in patients. The capacity to quantify X chromosome activity at chromosome-scale and patient resolution should also inform the study of female-specific gene regulation more broadly, an area that remains significantly underexplored relative to autosomal transcriptional programs. The interplay of escape, skew and variant exposure extends well beyond immunity into neurobiology, cancer and other contexts contributing to human phenotypic diversity. Together, these findings establish lineage-specific XCI escape, XCI skew and variant exposure as three measurable determinants of sex-biased immune gene dosage in health and autoimmune disease.

## MATERIALS AND METHODS

### Human Samples

Deidentified PBMCs were obtained from adult healthy donors and patients with RA and SS (Supplementary Table 2) from Cardinal Bioresearch ply ltd. An ethics exemption was granted by the Garvan Institute of Medical Research Human Research Ethics Committee (HREC) as samples were commercially collected and processed. Whole blood was also collected from adult healthy donors under ethics under the Royal Children’s Hospital (Melbourne) HREC, reference number 95179, and written informed consent was obtained prior to blood collection. PBMCs were prepared from whole blood by Ficoll-Paque centrifugation and cryopreserved in RPMI/10% DMSO/50% foetal calf serum (FCS). All research was conducted in accordance with The Code of Ethics of the World Medical Association (Declaration of Helsinki) for experiments involving humans.

### Mice

C57BL/6 JAusb (C57BL/6J) mice were purchased from Australian BioResources (ABR; Moss Vale, Australia). CAST/EiJ mice were kindly provided by Dr. Jacky Stoeckli. Mouse lines were maintained and managed at ABR and the Garvan Institute of Medical Research Biological Testing Facility (BTF). Following F1 intercross of C57BL/6J and CAST/EiJ mice, bone marrow was harvested by centrifugation of hind leg femur and tibial bones, prior to FACS and long-read sequencing. All experiments conformed to the current guidelines from the Australian Code of Practice for the Care and Use of Animals for Scientific Purposes. All mouse handling and experimental methods were performed in accordance with approved protocols of the Garvan Institute of Medical Research/St Vincent’s Hospital Animal Ethics Committee (AEC 25_21).

### Fluorescence activated cell sorting (FACS)

Thawed PBMCs were washed three times with RPMI/10% FCS, counted, then washed twice in PBS/10% FCS before resuspension in PBS/2% FCS through a 20mm filter. Immediately before analysis, DAPI was added and the cells were incubated for 5 minutes on ice. Sorting was performed on a FACS Aria III (BD Biosciences) using a single-cell gating strategy, with live cells selected based on DAPI exclusion using the V450 channel. Sorting was carried out at 4°C with a 70 mm nozzle. Cells were sorted into 1.5 mL Eppendorf tubes pre-coated with PBS/10% FCS to minimise cell adhesion, containing 100 µL of the same buffer. For samples DRG_0199_01, DRG_0198_01, DRG_1215_02, DRG_5467_02, DRG_2324_02 and DRG_2902_02 100,000 live single cells were sorted into a collection tube. For samples DRG_0175_01 and DRG_1409_01 40,000 live single cells were sorted into a collection tube. Cell numbers were chosen to match the targeted recovery for each capture approach.

### 10x Capture

Samples DRG_0199_01, DRG_0198_01, DRG_1215_02 DRG_5467_02, DRG_2324_02 and DRG_2902_02 were pooled together before capture by taking a 200ul aliquot from each FACS collection tube. This pool was captured on one channel of the 10x Chromium GEM-X platform using 5′ gene expression chemistry and targeting capture of 24,000 cells (4000 per sample). Samples DRG_0175_01 and DRG_1409_01 were captured using the 10x Chromium GEM-X On-Chip-Multiplexing platform with 5′ gene expression chemistry. They were pooled alongside two gorilla samples not included in this study, targeting 2,000 cells per sample. Samples were then identified via on-chip barcodes, removing the need for SNP-based demultiplexing. All captures used the Chromium Single-Cell 5′ Gel Beads and were performed by the Garvan Genomics Platform in the Garvan Institute of Medical Research. cDNA for all samples was produced using the 10x Genomics Chromium GEM-X Universal 5’ Gene Expression (vf3) assay. Library quality was assessed by quantification with a Qubit Fluorometer and fragment size verification with an Agilent TapeStation.

### WGS ONT Sequencing

Following PBMC thawing, 2×10^6^ cells per sample were used for ONT whole-genome sequencing. High molecular weight DNA was extracted using The Nanobind® PanDNA Kit following the manufacturer’s protocol. Libraries were prepared using the ONT Ligation Sequencing Kit (SQK-LSK114). Each sample was sequenced on a separate FLO-PRO114M flow cell using the ONT PromethION platform.

### Gene Expression ONT Sequencing

cDNA pools generated as described above were prepared using the ONT Ligation Sequencing Kit (SQK-LSK114) and were sequenced on FLO-PRO114M flow cells using the ONT PromethION platform. DRG_0341_01 and DRG_1409_01 were sequenced to a mean depth of 52,169 reads per cell and DRG_0199_01, DRG_0198_01, DRG_1215_02, DRG_5467_02, DRG_2324_02 and DRG_2902_02 were sequenced to a mean depth of 37,391 reads per cell.

### Processing of Nanopore Whole-Genome Sequencing Data

Human samples were sequenced at 30X depth on Oxford Nanopore PromethION flow cells. Basecalling was performed using the super accuracy model of Dorado [https://github.com/nanoporetech/dorado]. Reads were aligned to the human reference genome GRCH38 using minimap with the parameters -t48 -a -x map-ont. The resultant BAM was filtered for X chromosome reads using samtools, and then variants detected using Clair3^77^. The VCF was filtered for heterozygous, bi-allelic SNPs for input into scDaisyChain using bcftools^78^.

### Data Processing of 10x Nanopore single-cell data

Mouse 10x single-cell nanopore data was basecalled using the high accuracy model of Dorado [https://github.com/nanoporetech/dorado]. The fastq files were processed with the EPI2ME wf-single-cell pipeline [https://github.com/epi2me-labs/wf-single-cell] using the GRCm39 reference. Human 10x single-cell nanopore data was produced in two batches of pooled samples. The flowcells for each batch were basecalled using the super accuracy model of Dorado [https://github.com/nanoporetech/dorado], and processed using the EPI2ME wf-single cell pipeline [https://github.com/epi2me-labs/wf-single-cell] using the GRCh38 reference. The wf-single-cell workflow produced BAM files for each batch, where each read is tagged with the gene symbol / gene id (GN/GX), corrected cell barcode (CB) and corrected UMI (UB). In batch 1, which was performed using the 10x Chromium GEM-X On-Chip-Multiplexing platform, cell barcode to sample mapping is a priori known from the provided 10x barcode lists per channel. As the samples in batch 2 were pooled together and captured with 10x Chromium GEM-X platform using 5′ gene expression chemistry without on chip multiplexing, the cell barcode to sample mapping was obtained using Demuxafy^79^ with the VCF file produced from WGS above. Samtools was used to split the pooled BAMs into their respective individual samples using the CB tag^80^. Umi-tools dedup was used to deduplicate the tagged BAM with the options --per-cell and --per-gene^81^. The deduplicated BAM was used as input to scDaisyChain. The raw gene expression matrices produced by the wf-single-cell pipeline were also split by cell barcode for downstream analysis. Standard single-cell processing was performed using scanpy^82^. The demultiplexed raw gene expression counts matrix produced by the EPI2ME wf-single-cell pipeline was used as input. Cells with > 10% mtRNA content were filtered out, along with cells with less than 200 detected genes. Genes expressed in less than 3 cells were removed. Scrublet was used for doublet detection, and predicted doublets removed. The expression matrices were normalized to 10,000 reads per cell, and log1p normalized. Highly variable genes were determined using the built in scanpy function scanpy.pp.highly_variable_genes(). The highly variable genes were used for the PCA and UMAP generation. Celltypist was used for cell type annotation^83^.

### Variant Calling of C57/Bl6xCAST/EiJ

Paired end Illumina whole-genome sequencing reads for the parental inbred C57/Bl6 and CAST/EiJ lines were aligned to the GRCm39 reference genome using bowtie2^84^. Variants in both lines were called using GATK haplotype caller^85^. The CAST/EiJ VCF was filtered for variants that were not present in the C57/Bl6 line. All homozygous variants in the CAST/EiJ VCF were modified from a genotype call of 1/1 to 0/1 so that downstream software would treat them as heterozygous.

### scDaisyChain XCI pipeline

The scDaisyChain pipeline takes as input a coordinate-sorted and indexed BAM file containing GN, CB, UB tags for the gene name, cell barcode and UMI respectively, an unphased VCF containing heterozygous SNPs, a gene annotation GTF, and the original single-cell gene matrix directories. The pipeline consists of five main stages: allele-specific SNP counting, haplotype inference, read-level haplotype tagging, BAM splitting by inferred X-chromosome state, and generation of Xa/Xi gene and transcript matrices.

First, per-cell allele-specific SNP counts are generated from the input BAM and VCF. Only heterozygous, biallelic SNPs are considered. For each SNP, reads overlapping the variant position are extracted using pileup-based traversal of the BAM file. Secondary/supplementary alignments are not considered by default, and a minimum base quality score threshold for counting can be set (default 10). For each informative read, the cell barcode was taken from the CB tag and the gene assignment from the GN tag. Bases matching the reference allele, alternate allele, or neither allele are counted separately for each combination of cell barcode, SNP, and gene. This produced a SNP count table containing the cell barcode, genomic position, gene, reference and alternate alleles, reference count, alternate count, total count, and other-base count.

Where a GTF annotation is provided, SNP positions are additionally annotated against gene intervals. SNPs were first matched to exact GTF gene overlaps, then to genes within a user-defined flanking window, and finally to the nearest gene within a maximum distance threshold when no overlap or flanking match was found. The gene assigned by the read-level GN tag is compared with the GTF-derived gene annotation. Rows where the read-level gene assignment conflicted with the GTF annotation are optionally removed. SNP positions with evidence for multiple read-assigned genes and multiple nearby GTF genes could also be removed to avoid ambiguous gene assignment. When filtering removed SNP-count rows, the read IDs supporting those removed rows were recorded so that the same reads could optionally be excluded from downstream matrix construction. Haplotype inference was then performed from the allele-specific single-cell count table as described below by the scDaisyChain phasing algorithm in detail below.

After haplotype phasing, reads in the original BAM are tagged with haplotype and active/inactive-X information. For each read, all overlapping phased SNPs are identified from the phased VCF. At each informative SNP, the observed read base was compared with the allele assigned to haplotype 1 and haplotype 2. Haplotype support was summarised by the number of SNPs supporting each haplotype. Reads with stronger support for haplotype 1 were assigned to H1 and reads with stronger support for haplotype 2 were assigned to H2. The BAM was annotated with tags recording haplotype quality scores, haplotype SNP counts, the number of informative SNPs, and the final haplotype assignment. The read-level haplotype assignment was then combined with the cell-level active-X assignment produced during the scDaisyChain phasing step. For cells where X1 was inferred to be active, H1-supporting reads are labelled Xa and H2-supporting reads were labelled Xi. Conversely, for cells where X2 was inferred to be active, H2-supporting reads are labelled Xa and H1-supporting reads are labelled Xi. Reads without a cell barcode, reads from cells without an active-X assignment, and ambiguous reads are labelled separately. The tagged BAM are then split into separate BAM files corresponding to haplotype-specific reads and active/inactive-X reads, including X1, X2, Xa, Xi, ambiguous, and unknown categories. Each split BAM is indexed after writing.

Finally, gene count matrices are generated from the split BAMs. Gene-level matrices are generated using the GN tag. Reads could be excluded from matrix generation using the dropped-read ID list produced during the SNP-count filtering stage. The final output consisted of matched matrices for the X1, X2, Xa, Xi, ambiguous, and unknown read categories, enabling downstream quantification of allele-specific and active/inactive-X expression at gene or transcript resolution.

### Outline of the phasing scDaisyChain algorithm

Allele-specific single-cell count tables were used to infer chromosome-wide haplotypes using a weighted graph partitioning approach. Let *C* be the set of cells and *S* the set of heterozygous SNPs. For each SNP *s ε S*, let *s_R_* and *s_A_* denote the REF and ALT alleles. The input data are represented as an allele-by-cell count matrix *X*, where *X_a_*_,*c*_ is the number of reads in cell *c* supporting allele *a*.

Rows with missing or uninformative gene annotations were removed before phasing. For each SNP, total allele-informative coverage was calculated across all cells as

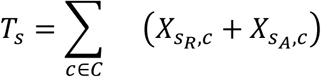

SNPs are retained if they pass a minimum read threshold and allele-balance filter:

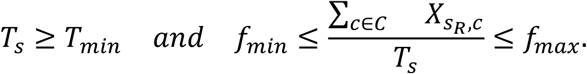

In the analyses described here, the default minimum total read threshold was (T_{\min}=10), and the minimum allele fraction threshold was (f_{\min}=0.01). This filtering step was used to retain SNPs with sufficient evidence for phasing while removing SNPs with very low support for one allele, which are less likely to represent informative heterozygous variants.

For graph construction, allele counts were converted to binary detection values indicating whether each allele was observed in each cell:

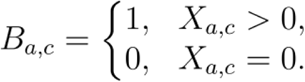

Each retained REF or ALT allele was represented as a graph node. Pairwise allele concordance was then calculated as a scaled co-detection score. For two allele nodes (a) and (b), the edge weight was defined as

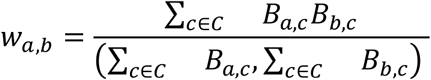

This weighting scheme assigns high weight to allele pairs that are consistently detected together across informative cells, while downweighting allele pairs with inconsistent co-detection. Direct edges between the REF and ALT alleles of the same SNP were removed before graph partitioning, because these alleles represent mutually exclusive alternatives at a single variant and should not directly attract one another during haplotype inference. The resulting weighted allele graph was partitioned into two communities using leading-eigenvector community detection implemented in iGraph, with edge weights included during partitioning. The two communities were interpreted as the two inferred haplotypes. Let *φ*(*a*) denote the community assigned to allele *a*. After each partitioning step, SNPs for which the REF and ALT alleles were assigned to the same community were considered inconsistent with a two-haplotype model. A SNP was retained after Stage 1 only if

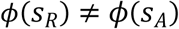

SNPs for which *φ*(*s_R_*) = *φ*(*s_A_*) are removed. The graph was then rebuilt using the remaining SNPs and repartitioned iteratively until no additional same-community REF/ALT SNP pairs remained. The final Stage 1 haplotypes were defined by the two graph communities:

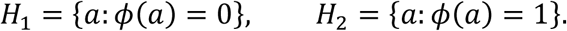

To assign the active X chromosome in each cell, counts from alleles assigned to *H*_1_ and *H*_2_ are aggregated across genes. The active-X state of cell *c* is defined as

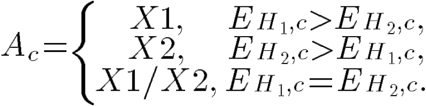

where *E*_*H*_1,*C*__ and *E*_*H*_2,*C*__ are the total haplotype-specific expression values in cell *c*. SNPs removed during Stage 1 were reconsidered in a Stage 2 fill-in step using the per-cell active-X assignments inferred from the high-confidence Stage 1 haplotypes. For a discarded SNP *s*, a cell was considered informative if it had an active-X assignment of either *X*1 or *X*2, and if the REF and ALT allele counts for that SNP were unequal:

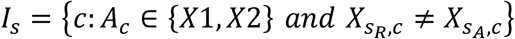

Each informative cell votes for one of the two possible haplotype orientations. ALT is assigned to *H*_1_ if it receives more concordant votes than REF:

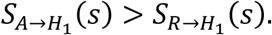

Conversely, REF is assigned to *H*_1_ if

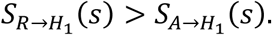

If both orientations receive equal support, the orientation is chosen at random. The final phased SNP set is the union of SNPs assigned by Stage 1 graph partitioning and Stage 2 active-X-based fill-in.

The final phased SNP set was defined as the union of SNPs assigned by Stage 1 graph partitioning and SNPs assigned by Stage 2 active-X-based fill-in. Final outputs included a per-SNP haplotype table containing REF and ALT alleles, inferred H1/H2 assignments, gene annotation, phasing stage, and total coverage; a per-cell table containing the active X call inferred from the high-confidence Stage 1 haplotypes; and a phased VCF containing the inferred haplotype orientation.

### Benchmarking of mouse nanopore data

The C57BL/6 × CAST/EiJ vcf generated from sequencing the parental lines was treated as the ground truth for bench marking purposes. scDaisyChain was run on the BAM file produced by EPI2ME wf-singlecell, requiring SNP coverage of at least 10 reads and minor allele fraction of 0.01. Phasing accuracy was evaluated by comparing the allele assigned to haplotype 2 (H2) with the known reference allele (REF) for each evaluated SNP. A SNP was classified as correctly phased when the H2 allele was identical to the REF allele. Accuracy was calculated as the proportion of evaluated SNPs that were correctly assigned. In order to investigate the effects of SNP misclassification on allele specific X chromosome expression, a corrected VCF where all misclassified SNPs were flipped was produced and used to tag the single cell BAM file. The X chromosome gene expression matrices resulting from the corrected and uncorrected BAM files were compared, by aggregating counts across genes and across cells and comparing the fraction of expression from the Xi, as well as the overall expression from the Xi, Xa, X1 and X2.

### Simulation of single-cell XCI escape data

We generated simulated allele-count datasets using the original mouse single-cell allelic expression data as the starting point. Rather than generating a fully synthetic count matrix from scratch, we used the observed per-cell, per-SNP allele-count table as the starting point. This preserves the empirical structure of the dataset, including the observed distribution of SNP coverage, gene expression, cell barcode representation, and SNP-to-gene annotations. We then applied controlled perturbations to this baseline dataset to model the effects of XCI skew, XCI escape, SNP removal, cell removal, and read removal on haplotype inference.

Unless a specific parameter was being tested, simulations were performed with 25% of genes simulated as escapees, 50:50 XCI skew, 80% SNP removal, no cell removal, and no read removal. Thus, each parameter sweep varied one feature of the dataset while the remaining simulation settings were held constant at this baseline.

To model reduced cell numbers, we randomly removed a specified percentage of cells from the baseline dataset. SNPs were defined by chrX position, and a specified percentage of SNPs was then randomly removed. The remaining SNPs defined the set of variants available for downstream simulated phasing. Genes with at least one retained SNP were identified, and a specified percentage of these genes was randomly selected as simulated escape genes.

Each retained gene was assigned a target inactive-X expression fraction. Genes selected as escapees were assigned a gene-specific Xi fraction drawn uniformly from 0.10 to 0.50. Non-escape genes were assigned a Xi fraction from 0-0.10. This allowed escape genes to contribute a controlled, gene-specific fraction of reads from the inactive X, while non-escape genes were simulated as predominantly subject to XCI.

To model reduced read depth, we applied binomial thinning to the observed total REF+ALT allele count for each SNP-by-cell observation. In this procedure, each original allele-informative read was randomly retained according to the specified read-retention probability. This generated lower-depth simulated datasets while preserving the empirical coverage structure of the original mouse data, such that high-coverage observations tended to remain relatively high coverage and low-coverage observations tended to remain low coverage.

We then assigned a simulated active-X state independently to each retained cell. The probability of assigning X1 as the active X was controlled by the X1 fraction parameter, while the probability of assigning X2 was one minus this value. This parameter determined the simulated XCI skew. For the baseline simulations, the X1 fraction was set to 0.5, corresponding to 50:50 XCI skew.

Simulated REF and ALT counts were generated using the convention that/X1 corresponds to the ALT allele and/X2 corresponds to the REF allele. For each SNP-by-cell observation, retained allele-informative reads were split into active-X-derived and inactive-X-derived reads according to the gene’s assigned target Xi fraction. In cells where X1 was simulated as active, ALT reads were treated as active-X reads and REF reads as inactive-X reads. In cells where X2 was simulated as active, REF reads were treated as active-X reads and ALT reads as inactive-X reads. The resulting simulated REF and ALT counts were written to the final allele-count table.

To assess phasing accuracy, scDaisyChain was run independently on each simulated allele-count dataset using the default minimum read threshold of 10 and minor allele-fraction cutoff of 0.01. Because the simulated haplotype truth was known, inferred haplotypes could be directly compared against the expected allele orientation. For each replicate, SNP-level accuracy was defined as the proportion of phased SNPs matching the simulated truth after orientation correction. Accuracy summaries were generated for each simulation replicate and then aggregated across iterations with the same parameter settings. The original number of expressed chrX genes was determined by identifying all chrX genes with at least 10 reads in the original unaltered gene expression matrix, and was used as the denominator when calculating the proportion of phased genes at each parameter combination. For parameter-sweep analyses, overall phasing accuracy and chrX gene recovery were plotted against the parameter being tested. Mean values across replicates were shown, with shaded regions representing standard deviations.

### Analysis of 1KGenomes SNP positions

The 1KGenomes39–40 VCF was filtered for female individuals and chromosome X variants only. The RNA coordinate of the closest heterozygous SNP to the transcriptional start site and transcriptional end site was quantified using a custom python script. Genes were considered classifiable at the coordinate of the first heterozygous SNP from the TSS or TES. To further estimate the capacity of long read sequencing to connect transcript termini to informative heterozygous SNPs, the X chromosome BAM files from eight donors were analysed using primary alignments and gene assignments from the GN tag. Read alignment blocks were mapped onto GRCh38 transcript models, and the transcript with the greatest exonic overlap was retained for each read–gene pair. The furthest aligned position from the transcript 5′ end was used as read reach, and gene-level median and 75th-percentile reach were calculated across donors. For each individual, distances from the TSS or TES to the nearest informative RNA SNP were compared with these gene-specific reach values. SNPs were considered reachable by Nanopore sequencing when their distance was within the median or 75th-percentile reach, and reachable by Illumina when within 150 bp. The fraction of genes with a reachable SNP was then calculated per individual.

### Comparison of read based phasing with Human Samples

Readbased phasing was performed using WhatsHap38 with the BAM file and VCF produced from nanopore WGS of each human sample. In order to minimize phase switch errors, the VCF was filtered for high quality SNPs by requiring a min QUAL of 20 and a min genotype quality of 10 using bcftools. The results of the read based phasing were compared with the scDaisyChain phasing for each sample. Concordance between scDaisyChain and WhatsHap was assessed using SNPs that were phased by both methods. Comparisons were performed within WhatsHap phase sets, as WhatsHap reports local phase blocks, whereas scDaisyChain assigns SNPs to chromosome-wide haplotypes.

For each WhatsHap phase set, we identified all shared phased heterozygous SNPs and compared their phased genotypes between the two methods. Because the labels assigned to the two haplotypes are arbitrary, scDaisyChain genotypes were compared to WhatsHap in both their original orientation and after flipping the scDaisyChain haplotype labels across the whole phase set. The orientation giving the highest number of matching SNPs was used for that phase set. Phase-set concordance was then calculated as the number of concordant SNPs after best orientation divided by the total number of shared SNPs in the phase set. Concordance across all SNPs was measured as well as average concordance per phase set. The impact of discordant SNPs was measured by flipping the discordant SNPs after best orientation alignment in the scDaisyChain vcf, and continuing with the scDaisyChain pipeline to quantify Xi fraction per gene for the original scDaisyChain vcf and the corrected version.

### CpG Methylation Analysis of Nanopore Whole Genome Sequencing

After basecalling and alignment as detailed above, the MM/ML modification tags were transferred from the basecalled reads to the genome aligned BAM file using modkit repair [https://github.com/nanoporetech/modkit]. Modkit pileup was used to summarise the modifications at CpGs and produce bedmethyl files. CpG methylation calls on chrX were annotated using BEDTools86. CpG island annotations were used to classify each CpG as overlapping a CpG island, a CpG shore (±2 kb from CpG islands excluding the island itself), or non-CpG island using the chrX CpG island annotation from UCSC87. CpGs were then intersected with promoter/TSS windows from the defined as ±1 kb around annotated TSSs in the GTF, and overlapping gene/transcript names were collected into a comma-separated annotation field. The final annotated bedMethyl file retained the original methylation columns and added CpG feature and TSS/gene annotation columns for downstream integration with expression and XCI escape classifications. As there are multiple transcription start sites per gene, for each sample, we identified the most expressed transcript for each gene from the transcript-level single-cell matrix, then filtered th1e methylation annotations so promoter CpGs were assigned only to that representative transcript/gene. This data was then merged with the gene-level escapee classification to compare the % of methylation at promoter regions in escapee, inactive and unclassified genes from scDaisyChain across all cells in each sample, with a threshold 10% of Xi expression required for escapee classification, and at least 20 cells with SNP informative expression for genes to be considered classified.

### Donor and Cell-Type-Resolved Inactive X (Xi) Quantification

For each donor, Xi- and Xa-assigned reads were linked to individual cells using cell-specific barcode information. Because individual genes are often sparsely expressed at the single-cell level, Xi- and Xa-assigned reads were subsequently aggregated within donor–cell-type–gene strata to generate pseudobulk estimates of gene-level Xi fractions while preserving the underlying single-cell haplotype assignments.

Analyses were restricted to cell types present at sufficient abundance across all donors to permit reliable donor-level allele-specific quantification: Helper T cells, Monocytes, Natural Killer (NK) cells, Cytotoxic T cells, Mucosal-Associated Invariant T (MAIT) cells, Memory B cells, Regulatory T cells, and Naïve B cells. Individual donor–cell-type–gene observations were retained if they contained at least five total allelic reads (Xi + Xa ≥ 5) and detectable allelic signal in at least three cells The raw Xi fraction was calculated as:

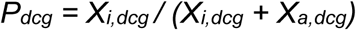

where X_i,dcg_ and X_a,dcg_ denote the total Xi- and Xa-assigned reads, and X_i,dcg_ + X_a,dcg_ is the total allelic count. The Xi expression fraction (p_hat_dcg_) was computed as follows:

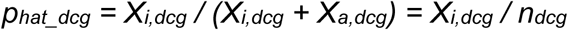

Raw Xi fractions were subsequently regularized using an empirical Bayes shrinkage framework. The allocation of allelic reads was modelled as:

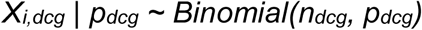

To capture the global distribution profile across the X chromosome, the latent parameter p_dcg was assigned a shared conjugate Beta prior:

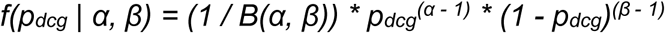

where B(·, ·) denotes the standard beta function. Under this generative model, the marginal distribution of the observed counts follows a Beta-Binomial architecture, a framework widely-used for single-cell allele-specific expression analyses^88^. The global hyperparameters α and β were estimated directly from the empirical distribution of raw data by maximizing the marginal log-likelihood across all globally retained baseline informative rows:

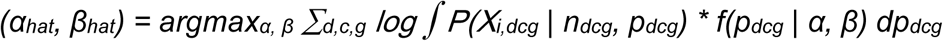

Utilizing the conjugate properties of the Beta prior, the posterior distribution of the latent inactivation fraction for each specific stratum given the observed allelic read counts updates analytically to a localized Beta distribution:

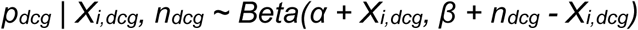

The posterior mean was used as the shrinkage-corrected estimate of the Xi fraction:

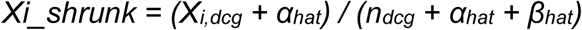

Unless otherwise stated, all downstream analyses used these shrinkage-corrected Xi fractions.

### Healthy Inactive X (Xi) Escape Atlas Construction

The baseline healthy Xi escape atlas was generated from healthy control donors (n = 4). For each gene and immune cell lineage, donor-level shrinkage-corrected Xi fractions were summarized across informative individuals. A gene was classified as escaping X-chromosome inactivation within a given lineage if Xi_shrunk_ >= 0.10 in at least two independent healthy donors16

Escaping genes were classified according to the number of immune cell lineages in which escape was observed: shared constitutive escape genes (loci escaping in >= 6 distinct immune cell lineages), intermediate escape genes (3 to 5 lineages), and selective or lineage-restricted escape genes (1 or 2 cell lineages). Loci failing to pass the escape threshold across all evaluated lineages were not classified as escape.

For targeted monocyte-versus-lymphoid lineage comparisons, the monocyte Xi fraction was contrasted directly against the arithmetic mean of the regularized Xi fractions across all mature lymphoid populations: ΔXi_lineage_ = Xi_monocyte_ - mean(Xi_lymphoid_). For visualization, loci with|ΔXi_lineage_| < 0.05 were considered neutral, whereas loci displaying absolute deviations of |ΔXi_lineage_| >= 0.10 were flagged as lineage-differential.

### scATAC-seq analysis

Cell type-resolved allele-specific chromatin accessibility on Xi and Xa was assessed using publicly available scATAC-seq data (ref. 30, https://www.10xgenomics.com/datasets/pbmc-from-a-healthy-donor-granulocytes-removed-through-cell-sorting-10-k-1-standard-2-0-0). For each gene, Xi/(Xi+Xa) accessibility ratios were computed using weighted.mean(), with allelic read counts used as weights to down-weight peaks with low allelic coverage. Cell types were mapped between the scDaisyChain and scLinaX nomenclatures, grouping Helper T and Regulatory T cells into CD4+ T, and Cytotoxic T and MAIT cells into CD8+ T categories.

For paired comparisons, Xi accessibility ratios were averaged across lymphoid (B, CD4+ T, CD8+ T, NK) and myeloid (Monocyte, DC) compartments per gene, and differences between compartments were assessed using a paired Wilcoxon signed-rank test (wilcox.test(paired = TRUE)), which accounts for gene-level correlation between compartments. For gene-level visualisation, chromatin accessibility tracks were rendered over a TSS-centred window (TSS ± 2,500 bp) using CPM-normalised merged bigWig files and the Gviz package. Allele-specific accessibility barplots were generated from the ref. 30 data for the same genes.

### Autoimmune Xi Analysis

Autoimmune analyses compared donor-level shrinkage-corrected Xi profiles between autoimmune patients and healthy controls. Rheumatoid arthritis (RA) donors (n = 3) were compared with healthy controls. Analyses required at least two informative donors per group. For each gene and immune lineage, group-level Xi values were calculated as the mean donor Xi fraction. Group differences were quantified as:

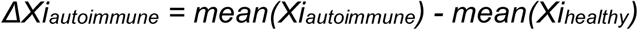

Cell-level UMAP coordinates were combined with donor, disease-status, cell-type and XCI-derived annotations. Cell-type-level Xi metrics, including escape fractions and disease-associated Xi shifts, were projected onto the embedding by assigning the corresponding value to all cells belonging to a given annotated cell type. Visualizations were generated in R using ggplot2 and displayed using continuous colour scales.

Principal component analysis (PCA) was performed on donor-level absolute Xi deviations from the healthy reference mean. Analyses were restricted to non-pseudoautosomal loci with |ΔXi_autoimmune_| >= 0.05, and features and scaled to unit variance prior to PCA. The primary visualization was generated using monocyte genes meeting the |ΔXi_autoimmune_| ≥ 0.05 criteria. A supplementary reference atlas including all expressed X-linked genes was generated without dysregulation filtering. Escape gains and losses were identified by comparing autoimmune and healthy donors using the same escape criterion (Xi.shrunk ≥ 0.10 in at least two independent donors). Statistical significance was assessed using 5,000 label permutations, and donor-level confidence intervals were estimated using 2,000 bootstrap resamples stratified by clinical group.

Genes meeting the reproducibility and effect-size thresholds were compared with a previously published monocyte trained-immunity signature associated with rheumatoid arthritis disease flare and synovial macrophage activation47. Analysis was restricted to X-linked genes expressed in our dataset. Overlap between the two gene sets was quantified using a two-sided Fisher’s exact test with the expressed X-linked gene set used as the background universe. Fold enrichment was calculated as the observed overlap divided by the overlap expected by chance. The significance of enrichment was assessed using Fisher’s exact p-values.

For the evaluation of the held-out Sjögren’s syndrome donor, a baseline healthy reference inactive X (Xi) value was established for each gene x cell-type combination using the control pool. For each individual Sjögren’s donor–gene–cell-type observation, the raw signed deviation was calculated as:

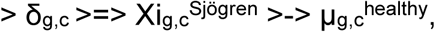

where μ is the healthy mean fraction, and absolute deviation was captured as |δ_(g,c)|. For each gene, the Autoimmune Deviation Score was defined as the mean of the two largest absolute cell-type deviations:

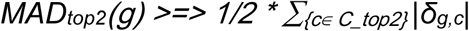

where C_top2 represents the subset of the two cell types displaying the largest absolute deviations for gene g.

### X-Chromosome Skew and Inactivation Dynamics

X-chromosome inactivation (XCI) skewing was quantified utilizing cell-level active-X chromosome haplotype assignments. For each individual donor or distinct donor–cell-type subgroup, the total pool of single cells assigned to the haplotype A (n_X1_) and the haplotype B (n_X2_) was explicitly counted. The raw H1/X1 active haplotype fraction was computed as follows:

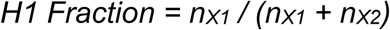

The final major-active-X fraction was defined as the maximum value of either the raw H1 fraction or its complement:

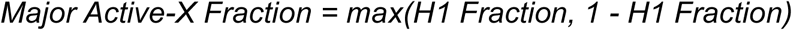

Values range from exactly 0.50 (balanced X-chromosome usage) to 1.00 (complete single-haplotype skewing). Donor-level confidence intervals were calculated using Wilson score intervals, and plotted as horizontal uncertainty segments ranked by effect magnitude. Lineage-specific cellular skew was evaluated by comparing each isolated donor–cell-type major active-X fraction directly against that donor’s overall major-active-X fraction:

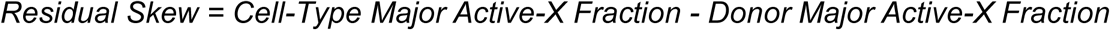

Positive values explicitly indicate increased skew relative to the donor baseline, whereas negative values indicate reduced skew. Lineage-level skew calculations required a minimum of 30 informative cells per donor–cell-type combination (min_cells_per_celltype = 30). Point sizes were scaled according to cell number.

### Variant Exposure, Genomic Integration, and Allelic Masking Models

Heterozygous X-linked variants from VCF records annotated using the Variant Effect Predictor (VEP) pipeline89 were integrated with donor-specific chromosome-scale haplotypes and lineage-specific active-X measurements. The haplotype carrying the alternative (ALT) allele was determined from phase annotations (H1_ALLELE_/H2_ALLELE_ or alt_on_H1/alt_on_H2). Variant Active Fraction was defined as the H1 active fraction when the ALT allele resided on H1 and 1 - H1 when the ALT allele resided on H2. Variants that could not be unambiguously assigned to a haplotype were excluded from downstream analyses. Variants were classified according to their Variant Active Fraction: (1) High exposure: Variant Active Fraction ≥ 0.70, (2) Low exposure: Variant Active Fraction ≤ 0.30 (3) Intermediate exposure: 0.30 < Variant Active Fraction < 0.70. Variant-level analyses required at least 50 informative cells spanning at least three immune cell types. Variants were further stratified into functional consequence categories including VEP HIGH-impact variants (nonsense, frameshift and essential splice-site variants), SpliceAI-disrupting variants (SpliceAI ≥ 0.10)90, splice-related variants, moderate-impact non-synonymous variants, and regulatory or untranslated region (UTR) variants.

### Statistical Implementations and Computational Reproducibility

All computational pipelines and statistical models were implemented using R (v4.3) and Python (v3.10) environments. Continuous associations were assessed using Pearson or Spearman correlation coefficients, as appropriate. Categorical overlaps and enrichment analyses were assessed using two-sided Fisher’s exact tests or hypergeometric tests. Multiple-testing correction was performed using the Benjamini–Hochberg false discovery rate (FDR) procedure. Unless otherwise stated, an FDR-adjusted P value < 0.05 was considered statistically significant.

## Supplementary Figures

**Supplementary Fig. 1:**
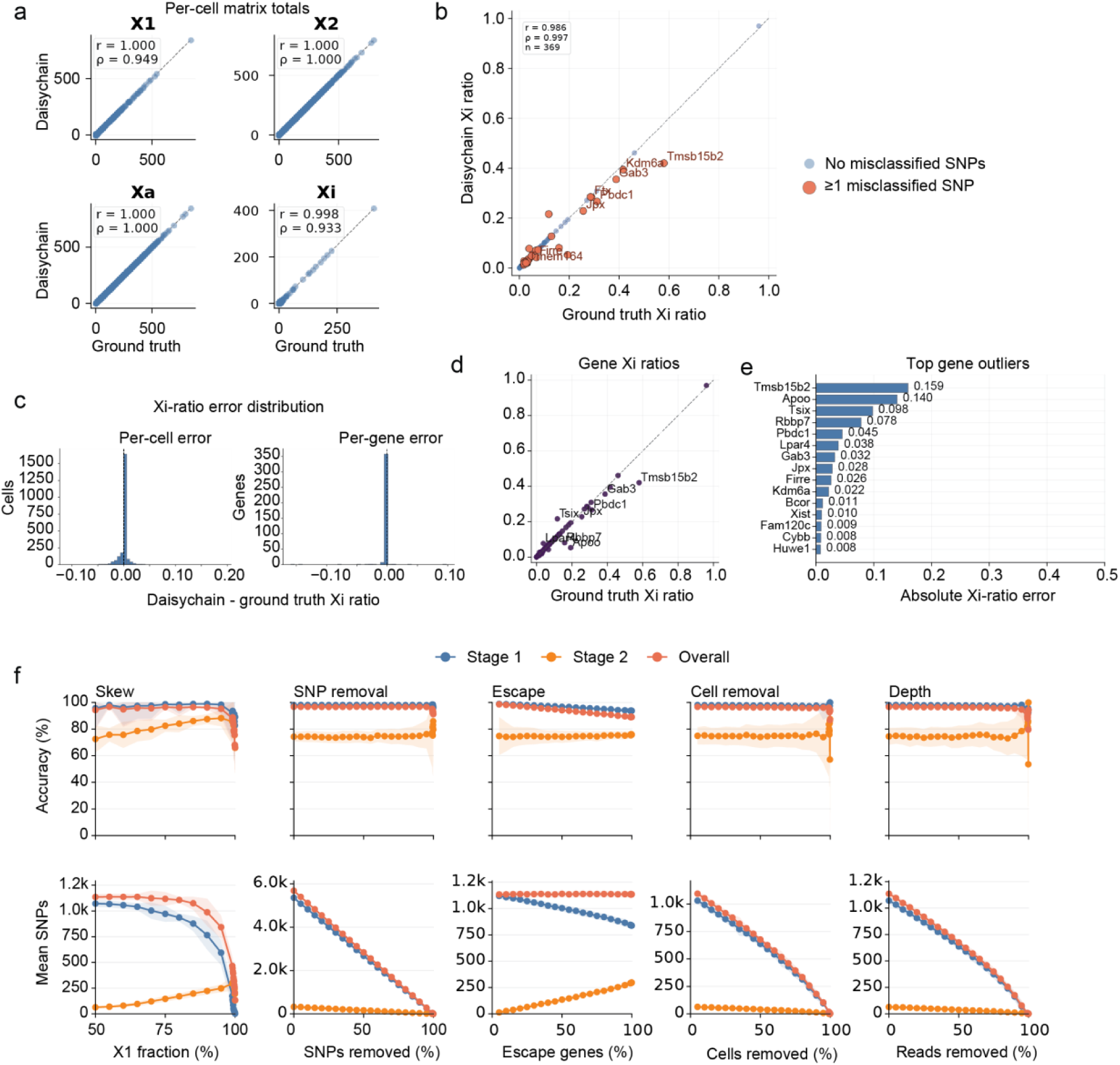
Validation of the scDaisychain algorithm in C57/BI6 X CAST/EiJ hybrid mouse model. **(a)** Total expression per cell from the two parental X haplotypes X1/X2 (top) and from the active and inactive X (bottom) in scDaisychain compared to the ground truth. **(b)** Fraction of expression from the inactive X per gene aggregated across all cells. Cells with at least one SNP misclassified by scDaisychain highlighted. **(c)** Fraction of expression from the inactive X per gene aggregated by cell, with genes with the highest Xi fraction deviation highlighted. **(d)** Distribution of the error in scDaisychain Xi fraction per cell and per gene compared to the ground truth. **(e)** Simulated single-cell allelic expression data using the C57/BI6 X CAST/EiJ mouse data as the starting point. XCI skew, percentage of SNPs removed and percentage of genes that escape XCI were set to 50%, 80% and 25% respectively. The percentage of the total number of cells (left) and percentage of the total number of allelic reads (right) in the dataset were varied with the other parameters fixed. **(f)** Stage 1, stage 2 and overall accuracy scDaisychain in C57/BI6 X CAST/EiJ simulations of XCI skew, SNP removal, Escape, Cell removal and Depth (top) with number of SNPs processed at each stage and overall (bottom).

**Supplementary Fig. 2:**
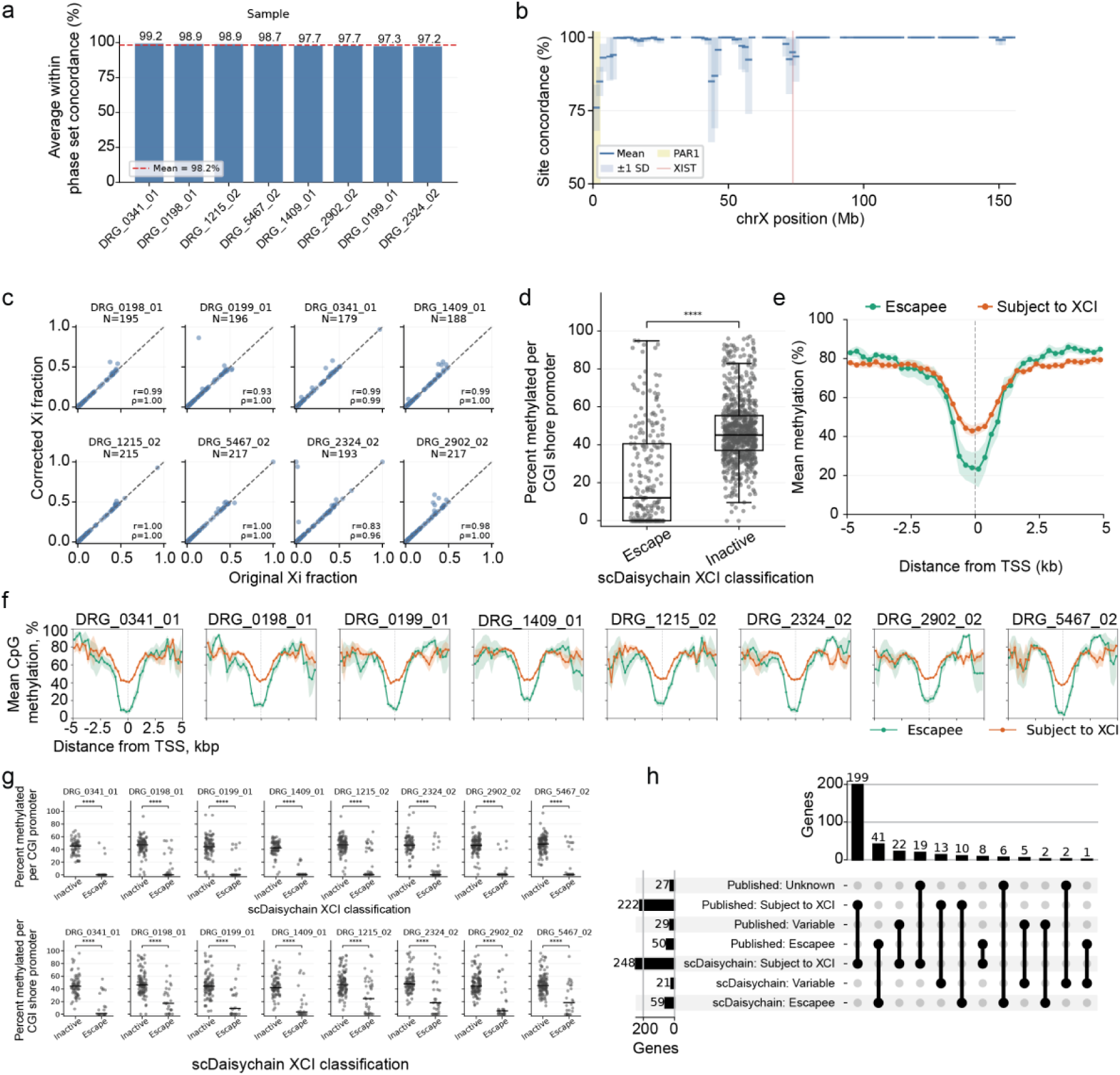
Validation of scDaisychain in human data. **(a)** Average phase set concordance accuracy per sample between scDaisychain and WhatsHap. Phase sets were identified with WhatsHap and accuracy is defined as the number of SNPs within a phase set that were concordant with the scDaisychain phase. As the H1/H2 orientation is arbitrary, both orientations are tested and the maximum is taken as the best orientation. **(b)** Within phase set concordance between scDaisychain and WhatsHap in 2.5mb windows with a step of 1.25 mb along the X chromosome, averaged across 8 donors with SD highlighted. **(c)** Correlation of scDaisychain original Xi fraction, and a corrected Xi fraction where the scDaisychain SNPs that were discordant with the best orientation WhatsHap result are flipped to match the assignment from WhatsHap. **(d)** Percent methylated per promoter overlapping CGI shores across **8** donors. Promoter regions were defined as ± 1 kb from the transcription start site of the most expressed transcript per gene in our dataset. **(e)** Mean methylation percent per gene in 250bp windows of CGI/CGI shores approaching the transcriptional start site, averaged across all 8 donors with SD shaded. **(f)** Mean methylation percent per gene in 250bp windows of CGI/CGI shores approaching the transcriptional start site per donor. **(g)** Mean methylation percent per CGI (top) and CGI shores (bottom) overlapping promoter regions per individual. **(h)** Concordance of scDaisychain XCI status classifications with previously published classifications. Genes were defined as escapees by scDaisychain if at least 10% of their expression across all cells in a donor was from the Xi across all informative donors, and subject to XCI if below 10% Xi expression for all informative donors, and variable if escapee in some donors but subject to XCI in others.

**Supplementary Fig. 3:**
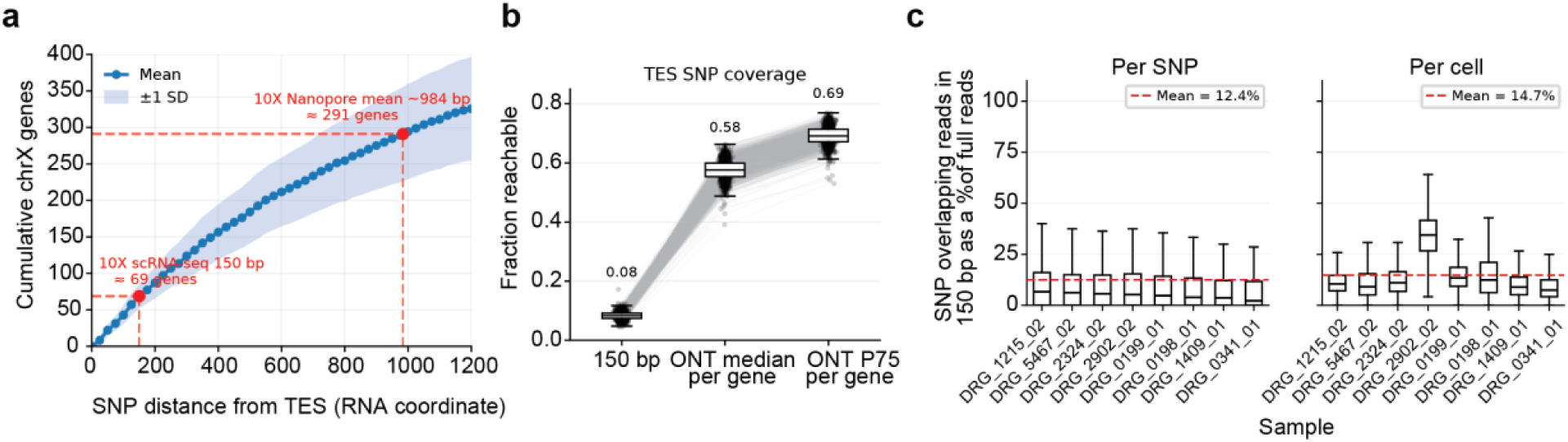
Comparison of scDaisychain with 10x Illumina single-cell sequencing. **(a)** Average number of genes with a heterozygous X chromosome SNP in female individuals in the 1kGenomes project ata range of distances from the transcriptional start site (TES), with typical 10x Illumina read length and mean read length in our 10x Nanopore human samples. **(b)** Average proportion of PBMC expressed X chromosome genes per female individual in the 1K genomes project with a SNP reachable in the last 150 exonic bp, using the median or 75th percentile read length of the gene from our 10x Nanopore human samples. **(c)** Percentage of SNP overlapping reads for each donor per SNP (left) and per cell (right) when truncating the reads to 150 bp compared to the full length reads.

**Supplementary Fig. 4:**
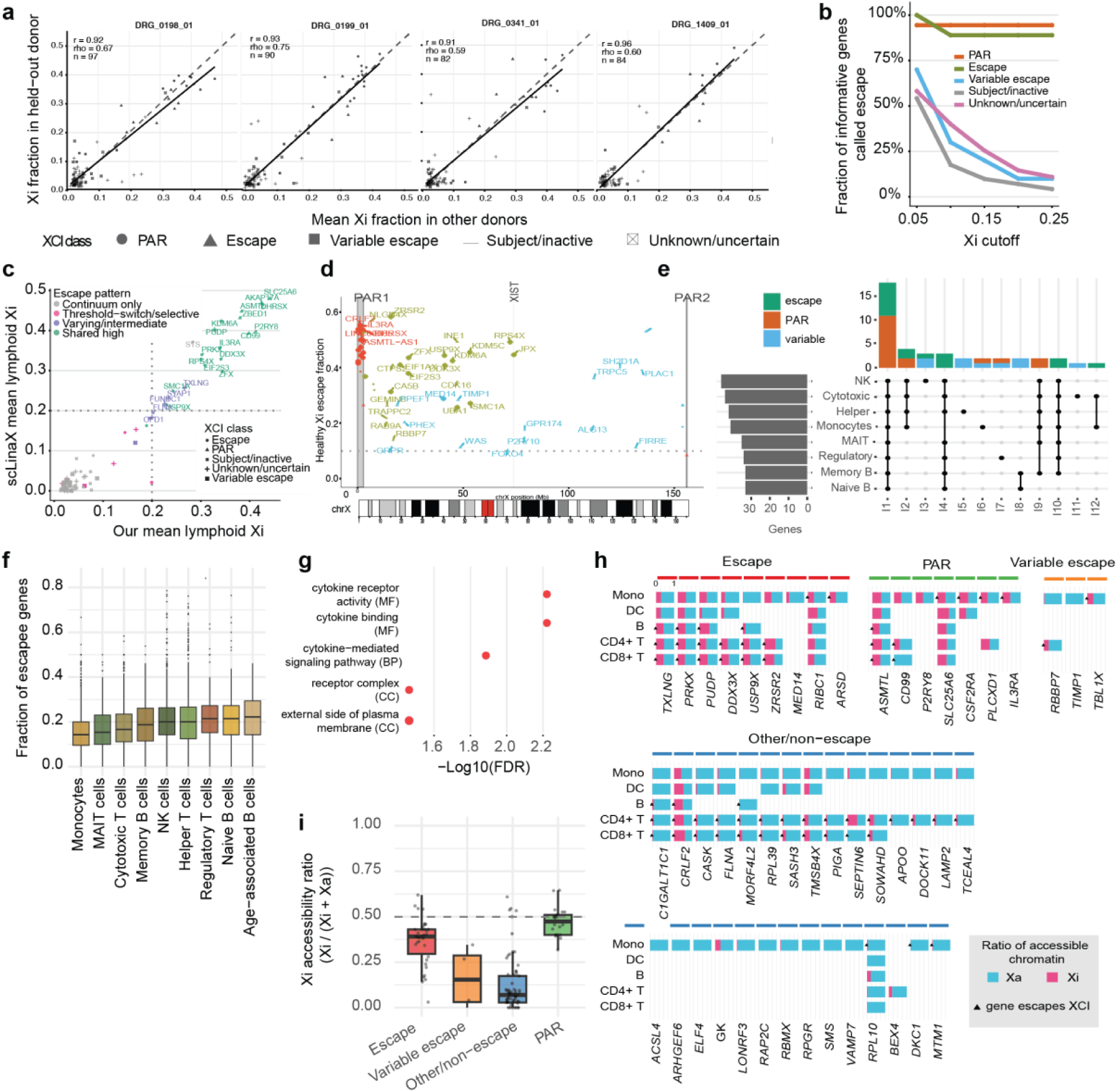
Characterisation and validation of X-chromosome inactivation (XCI) escape across human immune cell types. **(a)** Comparison of shrinkage-corrected XCI fractions for X-linked genes between individual donors and the mean across the remaining donors. Each point represents a gene, coloured according to its inferred XCI class. Pearson correlation coefficients are indicated for each comparison. **(b)** Sensitivity of XCI classification to the escape threshold. The proportion of genes retaining their original classification is shown across increasing XCI cut-offs for each XCI category. **(c)** Comparison of mean lymphoid XCI fractions estimated in this study with values reported by scLinaX. **(d)** Distribution of escape genes across the X chromosome. Only genes with mean XCI fraction >0.1 are shown. The location of XIST, PAR1 and PAR2 regions annotated with X chromosome ideogram are shown for reference. **(e)** UpSet plot summarising overlap of escape and PAR genes across immune cell types. Bars indicate the number of genes in each intersection, with connected dots denoting the corresponding cell-type combinations. **(f)** Box plot showing fraction of expressed genes classified as escape within each immune cell type **(g)** Functional enrichment analysis of the most variable XCI genes across immune cell types. Significantly enriched Gene Ontology terms shown in brackets. CC = cellular component; BP = biological process; MF = molecular function; FDR = false detection rate (Bonferroni-adjusted). **(h)** Cell-type-specific accessibility of escape genes. Heat map summarising chromatin accessibility at promoters of escape, variable escape, other/non-escape and PAR genes across immune cell populations. Colours indicate promoter accessibility on the active (Xa) and inactive (Xi) X chromosomes.

**Supplementary Fig. 5:**
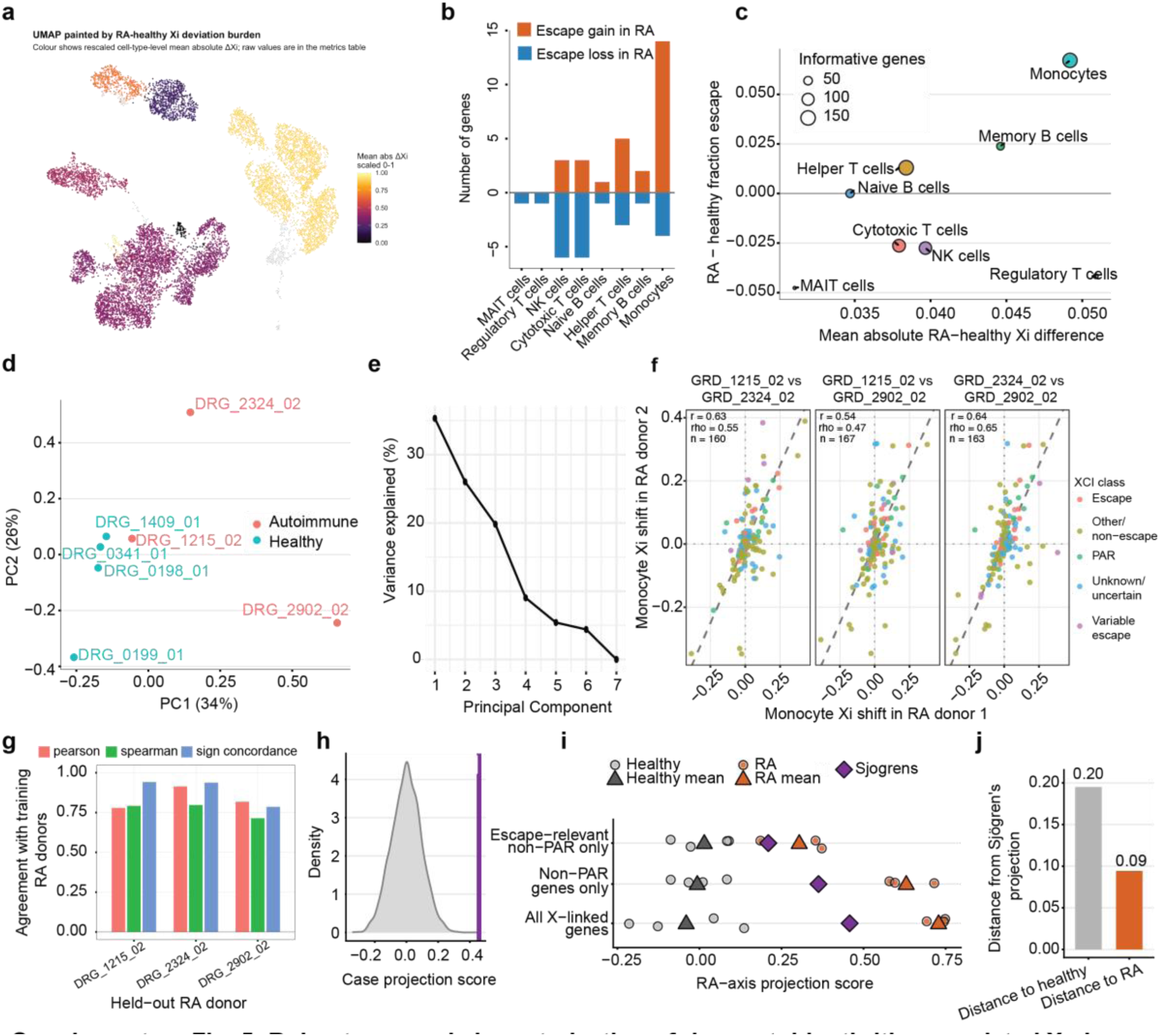
Robustness and characterisation of rheumatoid arthritis-associated X-chromosome inactivation (XCI) remodelling. **(a)** UMAP of immune cells coloured by the mean absolute difference in shrinkage-corrected XCI fraction (ΔXi) between healthy and rheumatoid arthritis (RA) donors for each cell type. **(b)** Number of genes exhibiting gain or loss of XCI escape in RA relative to healthy donors across immune cell types. Counts are restricted to genes informative in both healthy and RA donors for the corresponding cell type. **(c)** Summary of cell type-specific XCI remodelling in RA. The x-axis shows the mean absolute healthy-RA difference in XCI fraction, the y-axis shows the mean change in escape fraction (RA - healthy), and point size indicates the number of informative genes analysed. **(d)** Principal component analysis of XCI fractions in PBMCs across donors. Points represent individual donors coloured by disease status. **(e)** Scree plot showing the proportion of variance explained by each principal component in the monocyte XCI fraction matrix (from PCA in **Figure 4**). **(f)** Concordance of monocyte XCI shifts between pairwise combinations of RA donors. Each point represents a gene coloured according to its inferred XCI class. Pearson (p) and Spearman (rho) correlation coefficients are indicated for each comparison. **(g)** Leave-one-RA-donor-out stability analysis of the monocyte XCI signature. Bars show agreement between signatures derived from the remaining RA donors and the held-out donor measured by Pearson correlation, Spearman correlation and sign concordance. **(h)** Leave-one-RA-donor-out stability analysis of the monocyte XCI signature. Bars show agreement between signatures derived from the remaining RA donors and the held-out donor measured by Pearson correlation, Spearman correlation and sign concordance. **(i)** Projection of healthy, RA and Sjogren’s syndrome donors onto the RA-associated XCI axis using all X-linked genes, non-PAR genes only, or escape-relevant non-PAR genes. Circles denote individual donors and triangles/diamonds indicate group means. **(j)** Euclidean distance of the Sjogren’s syndrome donor from the healthy and RA group centroids in the RA-axis projection.

**Supplementary Fig. 6:**
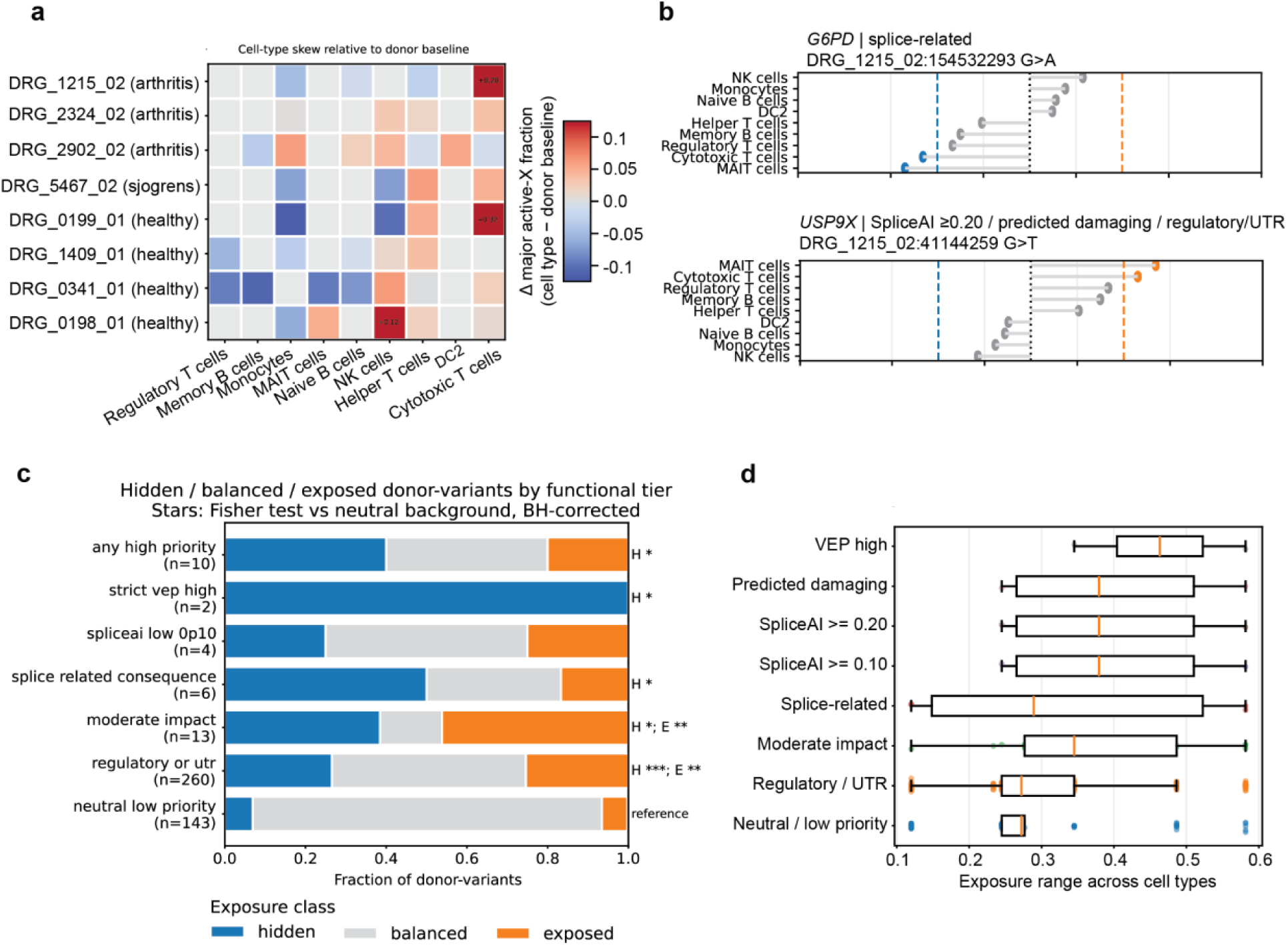
Cell-type variation in X-chromosome skew and consequences for X-linked variant exposure. **(a)** Heat map showing deviations in the active X-chromosome fraction for each immune cell type relative to the donor-specific baseline. Colours indicate the difference in active X fraction between each cell type and the corresponding donor average, highlighting lineage-specific variation in XCI skew. **(b)** Additional examples of cell-type-specific variation in variant exposure for representative X-linked variants. Points indicate the fraction of cells in which the variant-bearing haplotype is active for each cell type. Dashed blue and orange lines denote the thresholds used to classify hidden and exposed variants, respectively, and the dotted black line indicates balanced exposure. Annotated downloaded by Ensembl Variant Effect Predictor (VEP) and SpliceAl predictions. **(c)** Distribution of hidden, balanced and exposed donor-variant combinations across functional variant categories. Variant classes are defined using VEP annotation, SpliceAl predictions and predicted functional impact. Bars show the fraction of donor-variants assigned to each exposure class. Statistical significance relative to the neutral/low-priority background was assessed using Fisher’s exact test with Benjamini-Hochberg correction. H = hidden; E = exposed. * = p < 0.05; ** = p < 0.001; “* = p < 1×10-05. **(d)** Box plots displaying distribution of exposure range across immune cell types for donor-variant combinations grouped by functional annotation. Exposure range was calculated as the difference between the maximum and minimum exposure fractions observed across cell types for each donor-variant combination.

## Supplementary Tables

**Supplementary Table 1**: Single-cell statistics for human PBMC samples.

**Supplementary Table 2**: Donor information for human PBMC samples.

## Acknowledgements

We thank the members of the Weatheritt and Skvortsova laboratories for valuable discussions and critical feedback on the project and manuscript. We acknowledge the UNSW Restech HPC Scheme DOI: 10.26190/PMN5-7J50 for computational support and highlight that this research includes computations using the computational cluster Katana supported by Research Technology Services at UNSW Sydney DOI: 10.26190/669x-a286. We are grateful to Dr. Jacky Stoeckli for providing the CAST/EiJ mouse line. We also thank the Garvan Genomics Platform, particularly Dr. Chris O’Keefe and Dr. Alejandro Rios Villamil, and the Garvan Long Read Facility for technical assistance. Finally, we are grateful to the donors whose samples made this project possible.

## Author information

These authors contributed equally: Ksenia Skvortsova and Robert J. Weatheritt

## Author contributions

Conceptualisation: KS, RJW

Methodology: RJW, KS, CK, EMF, DK, HEK

Experimental design: KS, RJW, CK, EMF

Algorithm design: DK, with help from HK, RJW

Computational data analysis: RJW, DK, KS

Experimental work: AS, HGSV, EMF, KRK

Funding acquisition: RJW, KS, CK

Writing - original draft: RJW, KS, DK

Writing - review & editing: All authors

## Data and materials availability

The mouse and human single-cell long read RNA-seq raw sequencing data, and mouse WGS data generated for this study has been deposited at the European Nucleotide Archive (ENA) under accession number PRJEB115086. Simulated datasets, WhatsHap concordance benchmarking data, and CpG methylation benchmarking data are available at: https://figshare.com/s/49921651ec412cf584c2. Publicly available datasets and resources used in this study included 1000 Genomes Project variant call sets for chrX heterozygous SNP analyses39–40 and the Allen Human Immune Cell Atlas 41 (dataset DOI: 10.57785/ e9e1-wh09) for sex-biased immune-cell gene-expression analyses. Published X-chromosome inactivation annotations and/or estimates from Tukiainen et al.16 and Tomofuji et al.30 were used for comparison of Xi expression and/or chromatin accessibility, and the published rheumatoid-arthritis monocyte trained-immunity gene signature was used for enrichment analyses47.

## Code availability

The scDaisyChain package, usage steps, and analysis scripts for benchmarking are available on GitHub at https://github.com/weatheritt2/scDaisyChain and analysis scripts at https://github.com/weatheritt2/scDaisyChain_analysis_scripts

